# Stromal cells regulate malignant B-cell spatial organization, survival, and drug response in a new 3D model mimicking lymphoma tumor niche

**DOI:** 10.1101/2020.10.17.343657

**Authors:** Claire Lamaison, Simon Latour, Nelson Hélaine, Valérie Le Morvan, Céline Monvoisin, Isabelle Mahouche, Christelle Dussert, Elise Dessauge, Céline Pangault, Marine Seffals, Léa Broca-Brisson, Kévin Alessandri, Pierre Soubeyran, Frédéric Mourcin, Pierre Nassoy, Gaëlle Recher, Karin Tarte, Laurence Bresson-Bepoldin

## Abstract

Non-Hodgkin B-cell lymphomas (B-NHL) mainly develop within lymph nodes as densely packed aggregates of tumor cells and their surrounding microenvironment, creating a tumor niche specific to each lymphoma subtypes. Until now, *in vitro* preclinical models mimicking biomechanical forces, cellular microenvironment, and 3D organization of B lymphomas remain scarce while all these parameters constitute key determinants of lymphomagenesis and drug resistance. Using a microfluidic method based on the encapsulation of cells inside permeable, elastic, and hollow alginate microspheres, we developed a new tunable 3D-model incorporating extracellular matrix and/or stromal cells. Lymphoma B cells and stromal cells dynamically formed self-organized 3D spheroids, thus initiating a coevolution of these two cell types, reflecting their bidirectional crosstalk, and recapitulating the heterogeneity of B-NHL subtypes. In addition, this approach makes it suitable to assess in a relevant *in vitro* model the activity of new therapeutic agents in B-NHL.

## INTRODUCTION

Non-Hodgkin lymphoma (NHL) is a group of common hematological malignancies, with the majority of them originating from B cells. Follicular lymphoma (FL) and diffuse large B-cell lymphomas (DLBCL), the two most frequent B-cell NHL (B-NHL) (Smith et al., 2011; Teras et al., 2016), result from the malignant transformation of germinal center (GC) or post-GC B cells. FL are indolent lymphomas characterized by the occurrence of the t(14; 18)(q32;q21) translocation combined with additional recurrent somatic alterations (Huet et al., 2018). Two major subtypes of aggressive DLBCL, GC B-cell (GCB) and activated B-cell (ABC)-DLBCL, have been identified using gene expression profiling, reflecting their putative cell of origin and molecular alterations (Alizadeh et al., 2000). Despite a better understanding of the pathophysiology of these tumors, frontline therapy remains based on a combination of conventional chemotherapies such as CHOP (Cyclophosphamide, Hydroxydaunorubicin, Vincristine and Prednisone) and a monoclonal antibody against CD20. Nonetheless, 30-40% of DLBCL patients develop relapse or have refractory disease that cannot be cured with the standard immunochemotherapy (Sehn and Gascoyne, 2015). In addition, FL remains essentially an incurable disease and about 20% of patients progress or relapse in the first 2 years following treatment initiation, with a dismal prognosis (Huet et al., 2018).

It is now widely accepted that tumors constitute a complex ecosystem composed of many cell types regulated by biological, structural, chemical, and mechanical cues, that altogether participate in the effectiveness of therapeutic molecules (Bissell and Radisky, 2001). The tumor microenvironment of B-cell lymphomas contains highly variable numbers of immune cells, stromal cells, blood vessels, and extracellular matrix (ECM) and the interplay between these elements produces a tumor niche specific to each lymphoma subtypes (Scott and Gascoyne, 2014; Verdiere et al., 2018). FL exhibits a high dependence on a GC-like microenvironment where immune and stromal cells support survival, proliferation, and migration of malignant B cells. In turn, FL cells modulate the phenotype and function of their surrounding microenvironment. In particular, FL-infiltrating stromal cells are engaged in a bidirectional crosstalk with malignant B cells within infiltrated lymph nodes and bone marrow (Guilloton et al., 2012; Pandey et al., 2017). Conversely DLBCL have been proposed as less dependent on lymph node microenvironment but specific stromal signatures have been shown to impact DLBCL prognosis (Fowler et al., 2016; Scott and Gascoyne, 2014). Importantly, studies exploring functional interactions between lymphoma B cells and stromal cells have been essentially performed in 2D *in vitro* culture models whereas biomechanical forces and 3D organization emerged as key determinants of lymphoma pathogenesis and drug resistance.

*In vitro,* 2D culture of cell lines is the model mainly used to screen new molecules. However, this constitutes an “ideal” model where nutrients, oxygen, and drugs reach freely all cells. In addition, these cell cultures do not consider the spatial architecture of tissues, and lack microenvironment and ECM identified as tumor safety guards from chemotherapies. In this context, multicellular spheroids or tumor organoids represent promising models allowing high-throughput screening of anti­cancer drugs in versatile systems mixing several cell types and ECM components. While 3D culture models are increasingly developed for solid cancer (Clevers, 2016), their transfer to lymphoma modeling is still limited and includes: i) multicellular aggregates of lymphoma cells obtained using hanging drop method, which are useful for testing drug efficacy but do not account for the effect of cell-cell and cell-ECM interactions (Decaup et al., 2019; Gravelle et al., 2014), ii) 3D lymphoma organoid models integrating lymphoid-like stromal cells or integrin-specific binding peptides to recapitulate more accurately lymphoma microenvironment, but difficult to handle and low throughput (Tian et al., 2015).

In the present study, we aimed at providing a 3D lymphoma model that both integrates interactions between stromal cells and lymphoma B cells and is amenable to high throughput screening. Using a microfluidic method based on the encapsulation of cells inside permeable, elastic, and hollow alginate microspheres (Alessandri et al., 2013), we developped a new tunable 3D model incorporating ECM and/or stromal cells. We showed that lymphoma B cells and stromal cells form 3D spheroids whose internal architecture is driven by self-organization, thus initiating a bidirectional crosstalk and a co-evolution of these two cell types, recapitulating the heterogeneity of B-NHL subtype. In addition, this approach makes it possible to assess drug efficacy in a relevant *in vitro* B-NHL model.

## RESULTS

### 1 B-NHL cell lines form cohesive spheroids in scaffold-free alginate capsules

To produce spheroids from lymphoma B-cell lines, we used the cellular capsule technology, initially developed to generate spheroids from solid cancer tumor cells (Alessandri et al., 2013, 2016). Three B-NHL cell lines have been selected for encapsulation as representative models for GCB-DLBCL (SU-DHL-4), ABC-DLBCL (HLY1) and FL (DOHH2). From image-based analysis (N=656), the mean of the capsule diameter distribution was 215.9 ± 1.3 μm and the capsules contained an average of 18.2 ± 0.8 cells/ capsule (N=48) (Figure S1). During the first 11 days, SU-DHL-4 and HLY1 proliferated and aggregated to form spheroids. Alike the terminology used for 2D cultures, we called ‘3D confluence’, the stage when cells filled up the capsule and reached the internal wall of the alginate shell. Contrastingly, DOHH2 cells were not able to grow and form spheroids even after 17 days (Figure 1A) suggesting their requirement of a supportive microenvironment. Cell survival in SU-DHL-4 and HLY1 spheroids was evaluated at various culture time points using propidium iodide labelling. Upon spheroid dissociation and flow cytometry quantification, we found that the percentage of dead cells remained as low as 20­30% during the first week in culture and then increased to about 60% after 8 to 12 days (Figure 1B). As a corollary, EdU staining revealed that the early stages of spheroid growth were characterized by a high proliferation rate while post-confluent stages exhibited a significant decrease in cell proliferation concomitant with cell death increase (Figure S2). To determine whether this survival decrease was associated with the formation and growth of a dead cellular core, as observed in solid tumor spheroids (Däster et al., 2016; Sutherland, 1988), we performed immunostaining for cell proliferation (Ki67) and cell death (cleaved caspase 3). Besides an increased cell death after confluence (Figure 1B and Figure S2), neither Ki67 nor cleaved-caspase-3 regionalization was detected at prior or after 3D confluence (Figure 1C).

**Figure 1:**
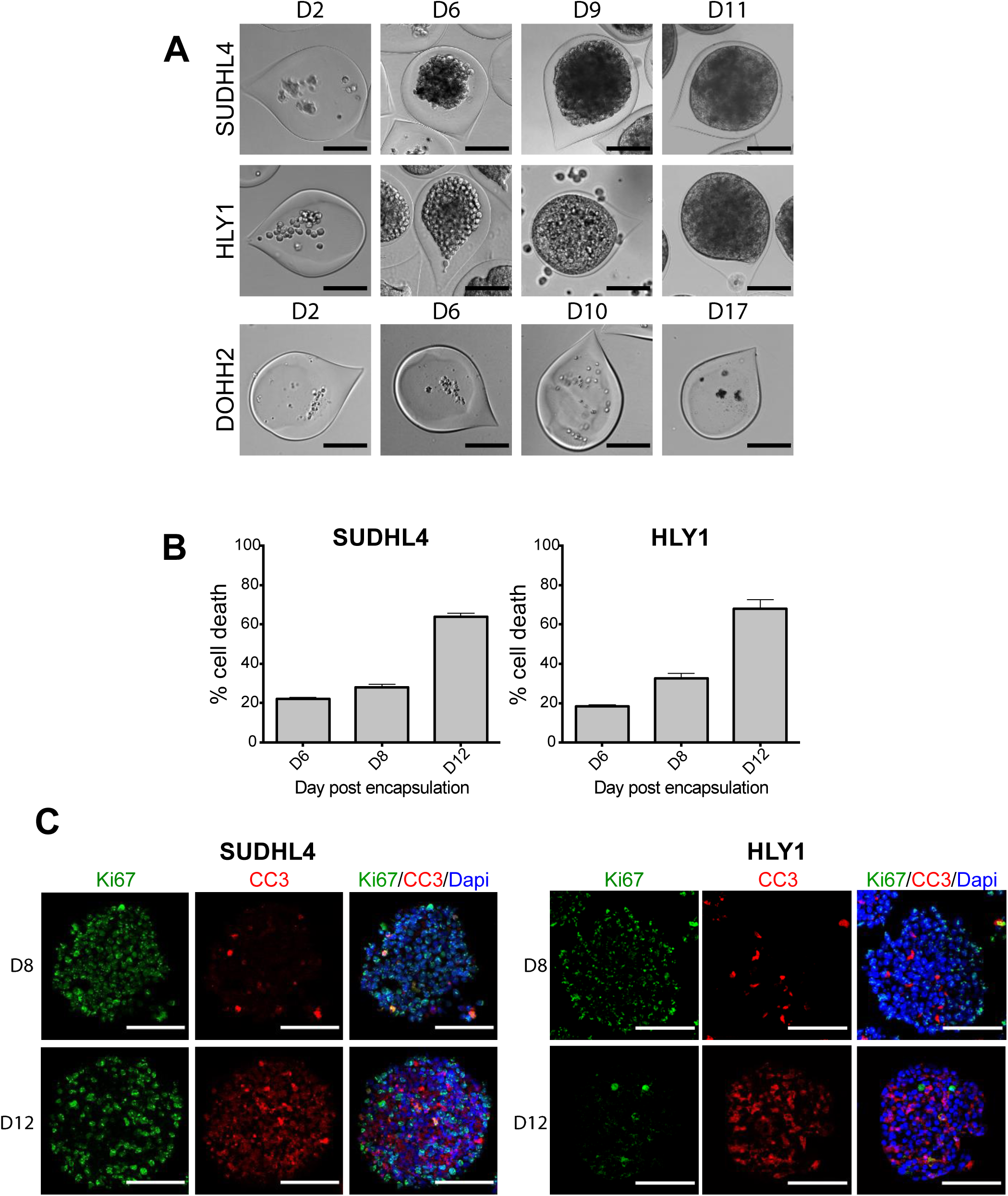
B NHL spheroids obtained by using the cellular capsule technology. **A:** Representative images of cell growth in alginate capsule shells over time for the various cell lines. Scale bar: 100μm. **B:** Percentage of SU-DHL-4 and HLY1 cell death measured after shell dissolution and cell dissociation at various culture time points. To evaluate the cell death propidium iodide (PI) (10 μg/mL) was added to the cell suspension and PI fluorescence was analyzed by flow cytometry (n=3). **C:** Immunostaining showing the repartition of proliferating and apoptotic cells in SU-DHL-4 and HLY1-derived spheroids at day 8 or 12 post-encapsulation. Five micrometers thick sections were stained with anti-Ki67 (Ki67, in green), and anti­cleaved caspase 3 (CC3, in red), to identify proliferating and dead cells, respectively. Nuclei are depicted in blue. Scale bar: 100μm.

To check that the capsule content was not only a confined cell suspension but formed truly cohesive spheroids, the alginate shell was dissolved in PBS containing 1 mM EGTA. As shown in figure 2A (and movie 1), the integrity of the cellular aggregates observed after confluence was maintained upon capsule dissolution, suggesting that 3D culture may induce the secretion of cohesive ECM. We thus compared by immunofluorescence the expression of laminin, fibronectin, and collagen I, three ECM components expressed in secondary lymphoid organs (Chen et al., 2012; Sobocinski et al., 2010), between cells cultured in suspension (2D) or in capsules (3D). SU-DHL-4 already expressed collagen 1 in 2D culture and upregulated laminin and fibronectin in 3D spheroids. Conversely, HLY1 displayed similar laminin expression in 2D and 3D cultures but increased fibronectin and collagen 1 in 3D spheroids (Figure 2B). Altogether these data argue for a promotion of ECM expression in 3D cultures of lymphoma cells.

**Figure 2:**
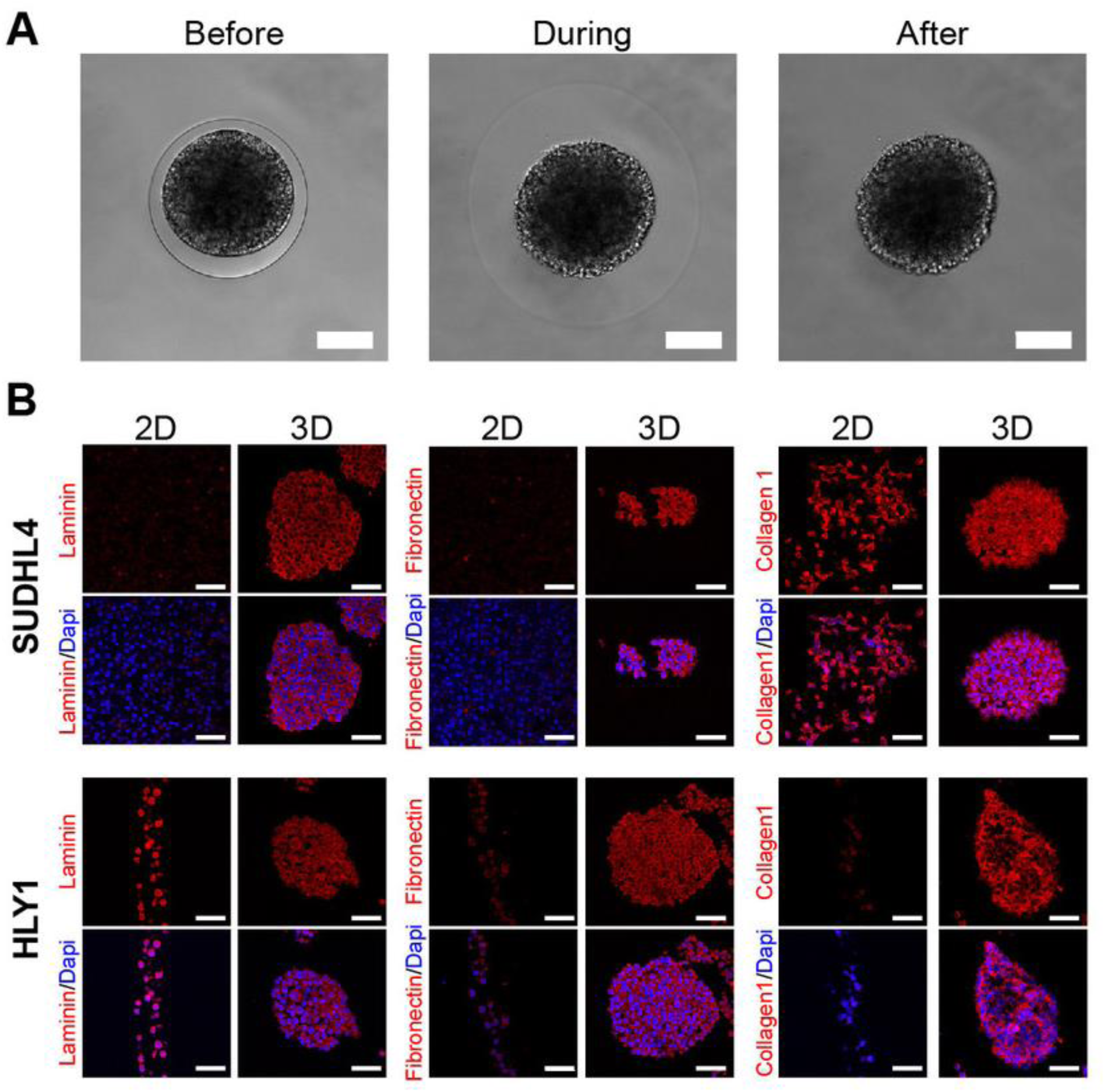
B NHL spheroids are cohesive and express extracellular matrix. **A:** Pictures showing the maintenance of SU-DHL-4 spheroids after the capsule dissolution by incubation in PBS containing 1 mM EGTA. Scale bar: 100μm. **B:** Immunostaining showing the expression of extracellular matrix in cells cultured in suspension (2D) or in spheroids (3D). After fixation, cell suspension was centrifuged and pellet was embedded in 2% agarose and then in paraffin, as performed for spheroids. Extracellular matrix components were visualized by staining with anti-pan laminin, anti-fibronectin and anti-collagen I antibodies on 5μm thick sections. Nuclei are depicted in blue. Scale bar: 50μm.

Altogether, these data show that the cellular capsule technology is a valuable approach to produce cohesive lymphoma spheroids from DLBCL.

### 2 Tumor B-cell and stromal cell co-encapsulation and 3D self-organization promote spheroid growth

Since B-NHL are organized as mixed aggregates of tumor cells and lymphoma-supportive non-malignant components (Verdière et al., 2018), we decided to aim for a more physiopathological model by introducing lymphoid stromal cells isolated from human tonsils (Tonsil Stromal Cells, TSC) (Ame-Thomas et al., 2007) and ECM components (Matrigel). We observed that lymphoma cell growth, evaluated by cell counting, was differentially regulated by the presence of ECM and stromal cells. Indeed, while the presence of stromal cells had no significant effect on HLY1 spheroid growth, they significantly increased the growth kinetic of SU-DHL-4 spheroids, and they even promoted DOHH2-derived spheroid formation and growth, which were not achieved in the absence of TSC (Figure 3A). These results suggest that our 3D model recapitulates the high TSC-dependence of FL B cells whereas DLBCL exhibit a more variable dependence (Ciavarella et al., 2018).

**Figure 3:**
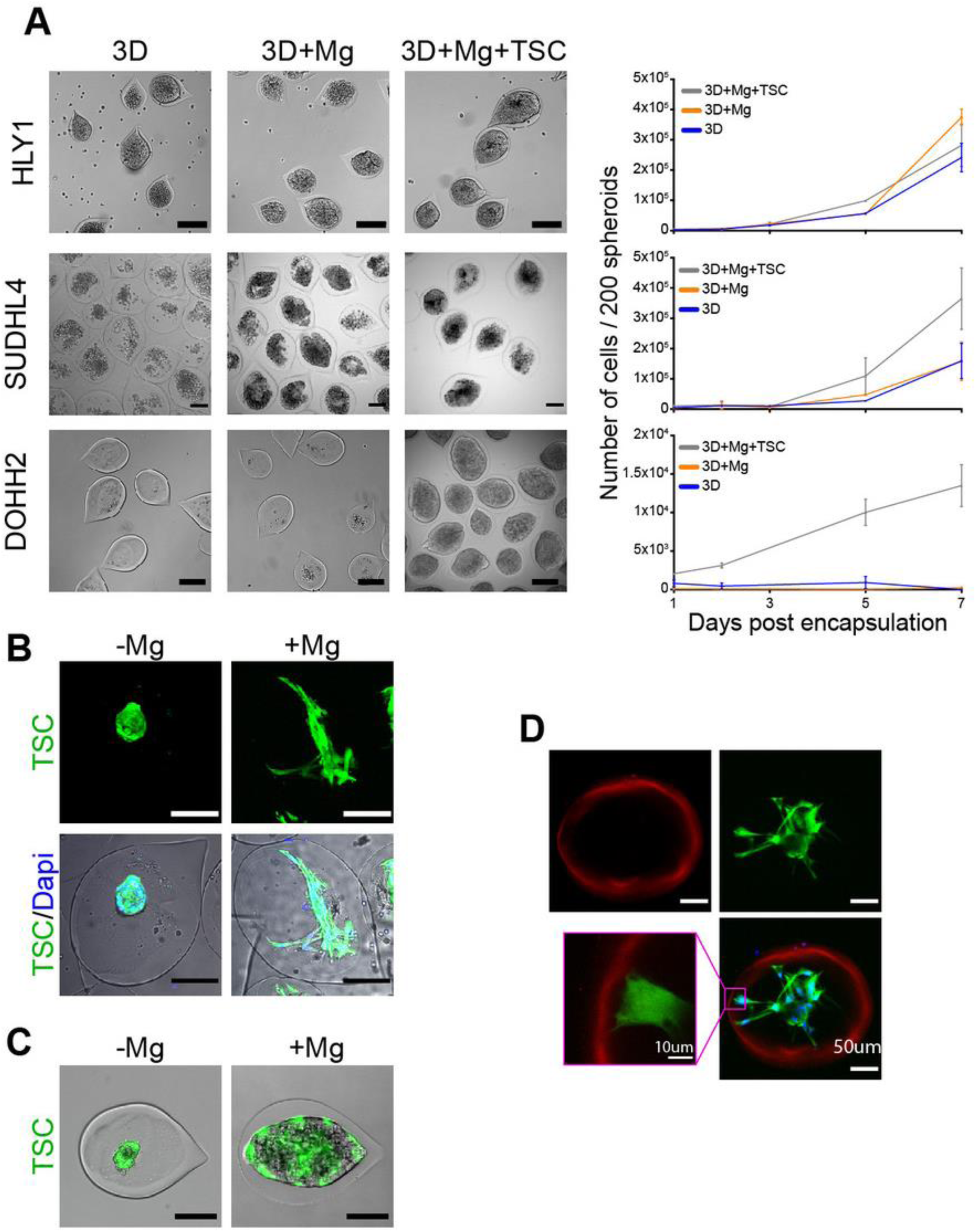
The establishment of a stromal network is necessary to reproduce stromal-dependent B NHL cell growth. **A:** Effect of Matrigel (Mg) and TSC on spheroid growth. Left panel, representative images of spheroids obtained from B cells encapsulated with or without Mg and TSC at D7 (HLY1), D6 (SU-DHL-4) or D10 (DOHH2) of culture. Scale bar: 200μm. Right panel, the cell number was evaluated at various culture time points in the different conditions (n=3). **B:** TSC network needs Mg to be established. *Left,* example of TSC encapsulated without Mg. *Right,* example of TSC encapsulated with Mg. TSC were visualized in green by stable expression of GFP and nuclei were stained in blue with Hoechst 33252. Capsules were imaged 3 days after encapsulation. Images were maximum intensity projection from z-stacks. Scale bar: 100μm. **C:** TSC network is necessary to induce DOHH2 spheroid growth. DOHH2 were co-encapsulated with TSC with or without Mg for 10 days. TSC were visualized in green by stable expression of GFP. **D:** TSC/Mg interaction. TSC were encapsulated in the presence of Mg. After 3 days in culture, capsules were fixed and immunofluorescence was performed. Mg and TSC were visualized by staining with anti-human pan laminin (in red) and anti-GFP (in green), respectively. Nuclei were counterstained in blue with Hoechst 33258. Images were maximum intensity projection from z-stacks. Scale bar: 50μm. Yellow square is a crop showing the hanging of TSC on Mg coating. Scale bar: 10μm.

We next sought to decipher how stromal cells play a supportive role on DOHH2 spheroid formation. DOHH2 cells were co-encapsulated with TSC in the presence or absence of a layer of Matrigel anchored to the inner wall of the capsules (Alessandri et al., 2016). In the absence of Matrigel, the capsule wall remained cell-repellent and stromal cells formed clusters inside the capsules but were unable to induce DOHH2 spheroid growth (Figure 3B and C). In contrast, in the presence of Matrigel, TSC spread onto ECM, leading to the formation of a ramified 3D network of stromal cells, that subsequently supported B cell proliferation (Figure 3D and movies 2-3).

We then monitored the stromal cell network dynamic by imaging capsules after 2, 24, 48, 72, and 96h of culture using confocal microscopy (Figure 4A). By solely focusing on stromal cells, we performed 3D+time image analysis and measured sphericity and branchness that both characterized the evolution of the stromal cell network complexity. High sphericity index indicated a compact morphology of the stromal network, thus offering a limited surface of interaction with B cells. Conversely, high branchness index was associated to a ramified network that increased the probability of B cell-stromal cell physical contacts. Stromal cell morphology was significantly modified as early as 24h after encapsulation, as revealed by the opposite variations of the sphericity and branchness indexes (Figure 4B and C). In addition, after a rapid initial change, both parameters underwent a slower but continuous evolution in time in the same correlated manner, indicating that the increased branching architecture of the stromal network was not a disorganized and randomly observed feature but was sustained by a self-organization process (Figure 4D).

**Figure 4:**
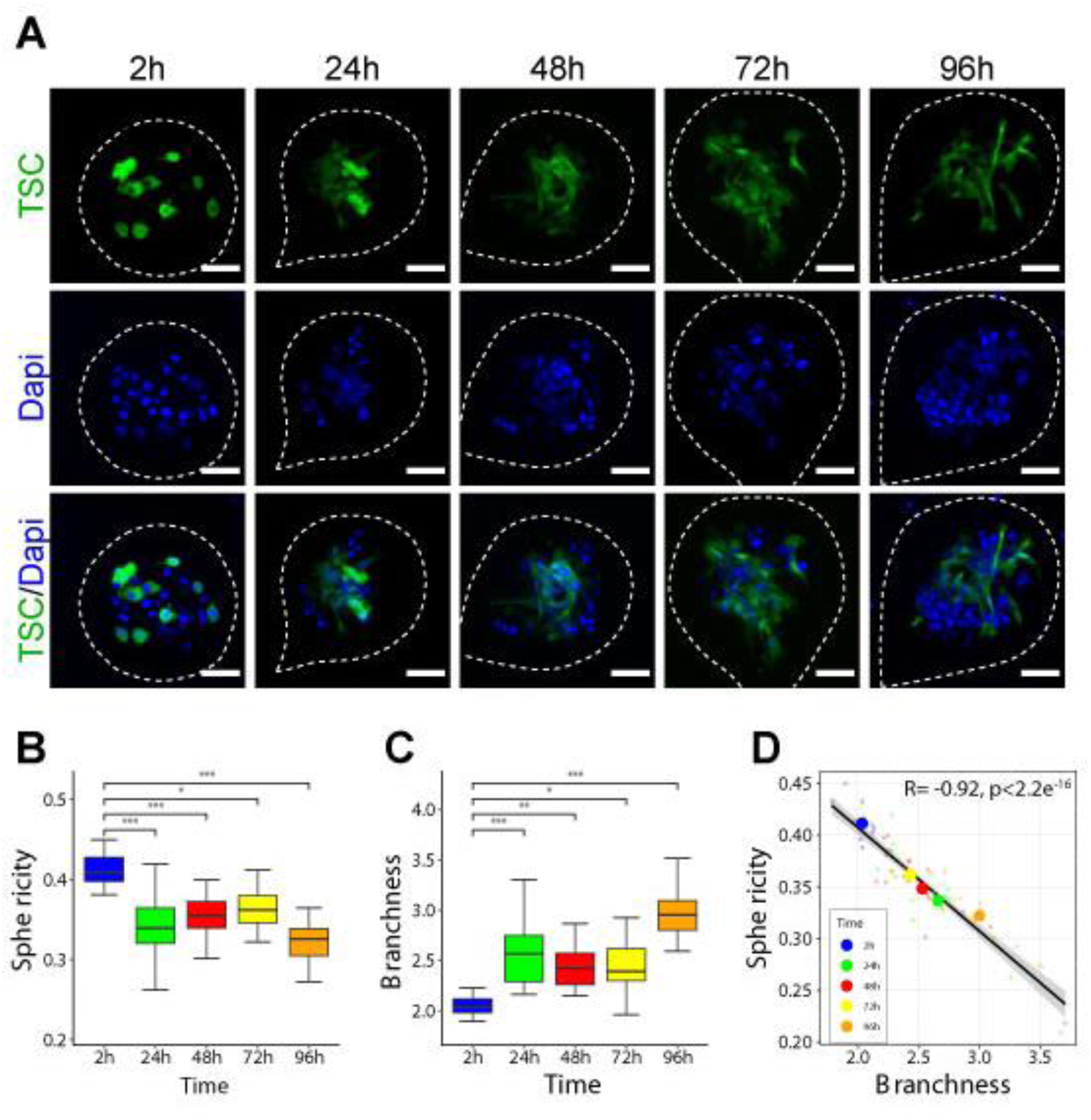
Self-organization of the stromal network. **A:** Representatives images of capsules containing DOHH2 with Mg and TSC at different culture time point. After fixation, 10 to 18 spheroids for each time point were placed in an Universlide (Alessandri et al 2017). Images shown are Z-projection. TSC are shown in green and all the analyses were performed on green signal exclusively. Scale bar: 50μm (**B**) and (**C**) Sphericity and branchness indexes of green particles contained in spheroids at different time after encapsulation, respectively. To determine significant differences, Anova was performed (*P<0.05, **P<0.01, ***P<0.001, ****P<0.0001). (**D**) Correlation of sphericity and branchness. For each time point, small dots represent individual spheroids whereas big dots represent the average values of all spheroids. Pearson correlation coefficient (R) was calculated using R software. The black line represents the regression line and the grey area represents the 95% confidence interval of the linear regression.

### 3 Microencapsulated 3D co-cultures recapitulate phenotypic features of B lymphomas

We decided to focus on the 3D TSC-DOHH2 co-culture to investigate the stromal/B-cell bidirectional crosstalk. Previously, we have demonstrated that FL B cells could contribute to the commitment of mesenchymal precursors into lymphoid­like differentiation (at least partly through the secretion of TNFα (TNF)) (Ame-Thomas et al., 2007; Guilloton et al., 2012). Interestingly, we revealed that 3D cell co­cultures induced a stromal cell phenotype similar to that observed *in situ* in FL lymph nodes (Figure 5). More specifically DOHH2 induced podoplanin (PDPN) expression in stromal cells while this was not the case in 2D co-cultures in which only the treatment of TSC cells with a combination of TNF and lymphotoxin α1β2 (LT), the two factors involved in lymphoid stroma differentiation, was able to trigger stromal PDPN expression (Figure 5 and Figure S3). Within spheroids, stromal cells thus acquired characteristics mimicking those observed *in situ*, suggesting that 3D spatial organization is an added value to recapitulate the pathophysiology of lymphoma tissues.

**Figure 5:**
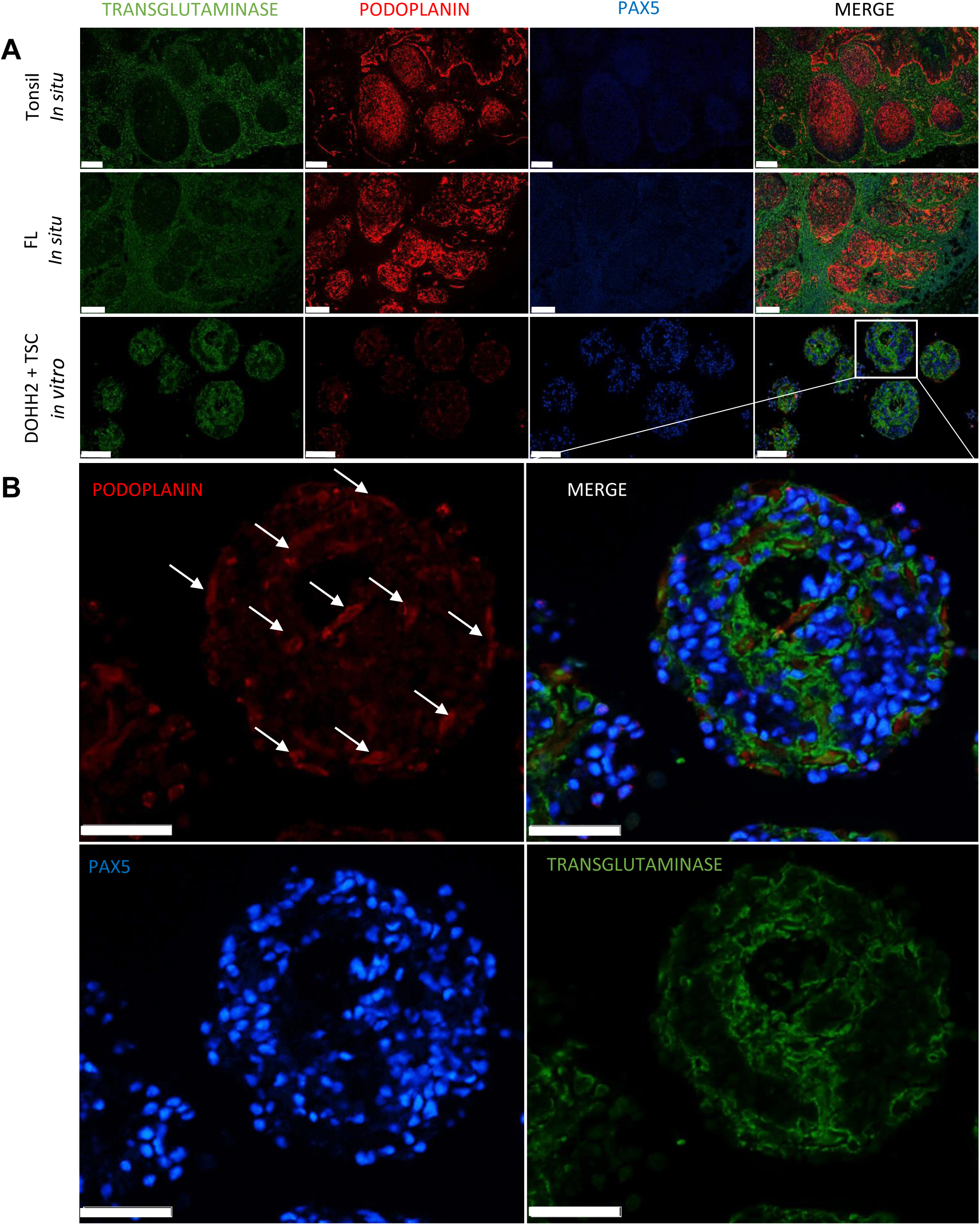
Spheroids recapitulate the FL stromal cell phenotype. **A:** Tonsil B cells (revealed with anti-PAX5 antibody in blue) interact mainly with stromal cells expressing podoplanin (red). In FL lymph node, tumor B cells interact with stromal cells expressing podoplanin (red) and transglutaminase (green) markers. Scale bar: 250μm. In the 3D model TSCs adopt a particular phenotype in contact with DOHH2 by expressing transglutaminase (in green) and podoplanin (in red) markers. Scale bar: 100μm. **B:** 3D model at a higher magnification represents DOHH2 (in blue) in close contact with the podoplanin (shown by arrows in red) and transglutaminase (in green) expressing TSC network. Scale bars: 50μm. All images were stained with multiplex immunofluorescence microscopy method and visualized by Nanozoomer software.

We next studied the gene expression profile (GEP) of DOHH2 cells cultured for 10 days either in monoculture in 2D (noted 2D), or in co-culture with TSC in 2D or 3D (respectively noted 2D + TSC and 3D + TSC). Data were then compared with the GEP of purified FL B cells and normal tonsil centrocytes (CC) obtained by Affymetrix microarrays. Expression of 37 genes involved in several transduction pathways including NF-κB cascade, migration signaling, cell cycle/quiescence, and apoptosis, was studied by RT-qPCR (Table 1, Figure S4). Analyses revealed a shift in GEP of B cells cultured in 3D compared to cells cultured in 2D with or without TSC (Figure 6 A). In particular, we observed that the expression of *CDKN1A* and *CDKN1B,* two regulators of cell quiescence (Heldt et al., 2018; Sgambato et al., 2000; Zhang et al., 2013), was similarly up regulated in 3D vs 2D co-cultures and in FL-B cells vs GC B-cells (Figure 6B). Moreover, *BCL2* was upregulated in DOHH2 after the 3D co-culture and was highly expressed in FL B cells (Figure 6C).

**Figure 6:**
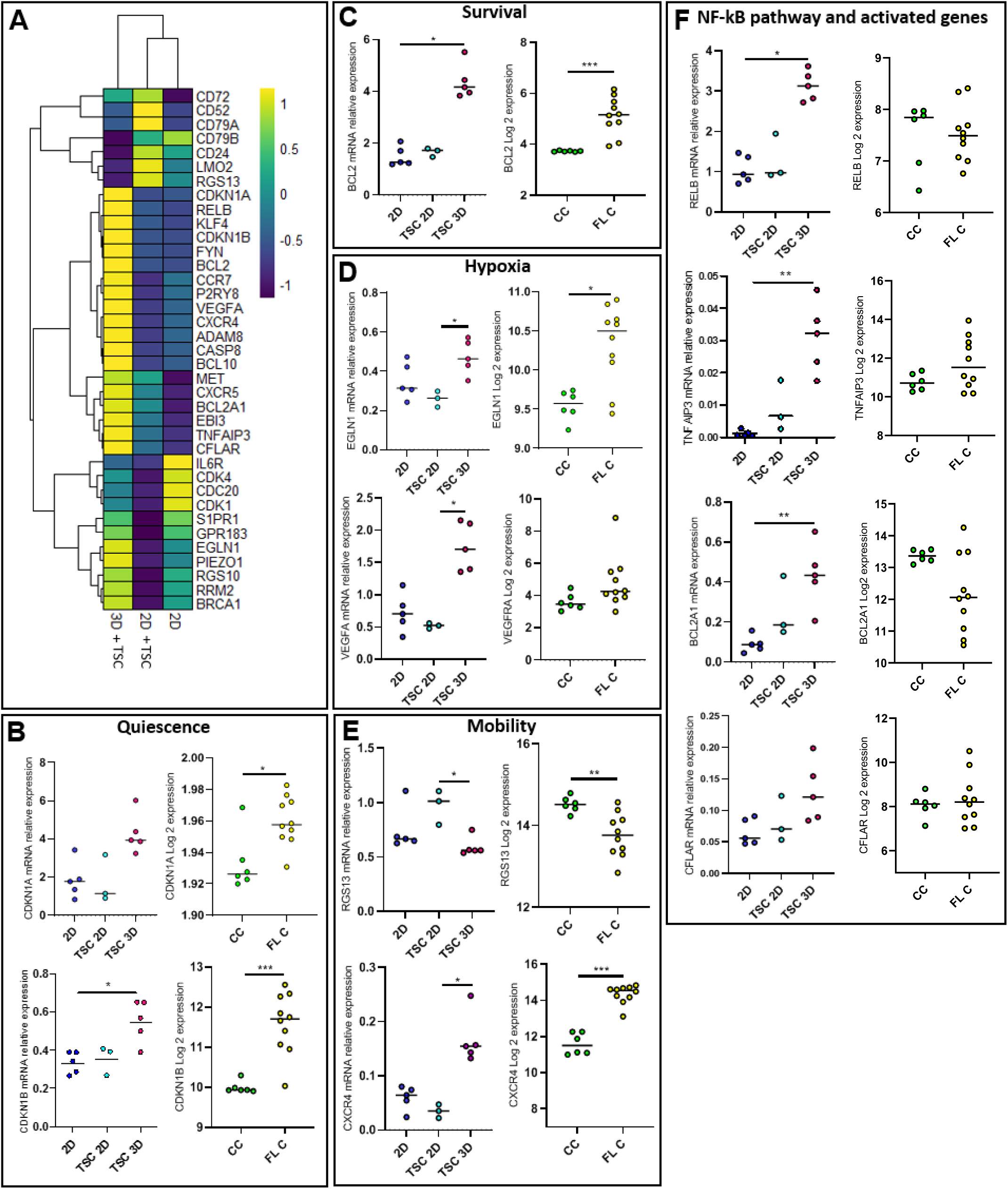
Comparison of DOHH2 and FL B cell gene transcriptional expression profile. **A:** Heatmap of DOHH2 gene expression levels evaluated by Real Time quantitative PCR (RT-qPCR) in cells cultured in 2D without (n=5) or with TSC (n=3), and in 3D with TSC (n=5). Analysis of DOHH2 transcriptional gene expression profiles, by RT-qPCR, for **B:** *CDKN1A* and *CDKN1B,* **C:** *BCL2,* **D:** *EGLN1* and *VEGFA,* **F:** *RGS13* and *CXCR4,* **G:** *RELB, TNFAIP3, BCL2A1* and *CFLAR.* DOHH2 were cultured in 2D without (n=5) or with TSC (n=3) and in 3D with TSC (n=5) for 10 days. Variation gene expression study between tonsil centrocytes (n=6) and FL lymph nodes tumoral Β cells (n=10) was studied by using Affymetrics micro-array analysis. A Kruskal Wallis test was performed for *in vitro* experiments and a two tailed Mann Whitney test was performed for the variation gene expression study between tonsil centrocytes (CC) and LF lymph nodes tumoral Β cells (FL C). (p<0.05*, p<0,01**). Bar represent the geometric mean of independent experiments.

**Table 1:** List of genes

| **Pathway** | **Gene** |
| --- | --- |
| **Cell survival** | *CFLAR*<br>*BCL2*<br>*BRCA1*<br>*BCL2A1* |
| **Hypoxia** | *VEGFA*<br>*EGLN1* |
| **B cell and BCR activation** | *EBI3*<br>*FYN*<br>*CD79b*<br>*CD79a*<br>*RRM2* |
| **Proliferation and differentiation** | *CD24*<br>*MET*<br>*CD72* |
| **Cell cycle** | *CDC20*<br>*CDK1*<br>*CDK4* |
| **Mechanosensitive** | *PIEZO 1* |
| **NFκB** | *TNFAIP3*<br>*BCL10*<br>*RELB* |
| **Quiescence** | *CDKN1B*<br>*CDKN1A* |
| **Confinement and motility** | *CXCR4*<br>*RGS13*<br>*P2RY8*<br>*S1PR1*<br>*CCR7*<br>*GPR183*<br>*CXCR5* |
| **Apoptosis** | *CASP8* |
| **Cell and matrix interactions** | *ADAM8* |
| **Adhesion** | *CD52* |
| **Others** | *RGS10*<br>*KLF4*<br>*LMO2*<br>*IL6R* |
| **Housekeeping genes** | *GAPDH*<br>*ABL1* |

The hypoxic tumor environment has been often reproduced in 3D tumor *in vitro* models (Benien and Swami, 2014). We confirmed this observation in our lymphoma 3D model, since both the upstream HIF1A transduction pathway regulator *(EGLN1)* and some HIF1A target genes *(VEGF, CXCR4)* were upregulated compared with 2D co-culture condition, reflecting, at least in part, the expression pattern obtained in FL B cells compared with normal CC (Figure 6D and E).

FL is a disseminated disease and we thus sought to determine the impact of 3D architecture on the expression of genes involved in cell confinement or cell migration. Interestingly, *RGS13,* known to limit GC size and GC B-cell number by interfering with CXCL12 and CXCL13 signals (Hwang et al., 2013), was specifically downregulated in 3D co-cultures and FL B cells. Conversely, *CXCR4* expression was similarly upregulated in FL-spheroids and FL B cells (Figure 6E).

Activation of the NF-κB pathway has been extensively described in B-NHL (Kennedy and Klein, 2018), leading us to investigate the expression of this pathway in our 3D model. Our analyses revealed an upregulation of the noncanonical *(RELB)* NF-κΒ pathway in spheroids compared to 2D co-cultures. Concomitantly, the expression of *TNFAIP3,* a negative regulator or the canonical NF-κB pathway, as well as that of *BCL21A* and *CFLAR,* two target genes of NF-κΒ, were increased in 3D compared to 2D co-cultures (Figure 6F). These results suggest that a functional non­canonical NF-κΒ pathway is activated in FL-spheroids.

Collectively, these results suggest that the co-culture of DOHH2 with TSC in 3D exhibits high degree of similarity with FL biopsies and partly mimics the lymphoma niche.

### 4 B-NHL sensitivity to chemotherapy in 3D vs 2D cell culture

We wondered whether the 3D architecture could alter the drug response of lymphoma Β cells. For that purpose, we evaluated the effect of doxorubicin, one of the components of the classical polychemotherapy used in Β-NHL (CHOP), on the cell death of Β-NHL cell lines cultured in 2D or 3D in the absence or presence of stromal cells. As shown in Figure 7A, we observed that the induction of apoptosis by doxorubicin tested at a concentration corresponding to its EC50 for the different cell lines (Figure S5), was abolished or significantly decreased in cells cultured in 3D compared to 2D. Based on our results showing that *BCL2* expression was upregulated in FL spheroids, we then tested the effect of a BCL2 inhibitor. ABT-199 induced massive cell death in DOHH2 co-cultured in 2D with or without stromal cells, as well as in 3D co-cultures, suggesting that malignant Β cells retained their dependence on BCL2 expression for survival (Figure 7Β).

**Figure 7:**
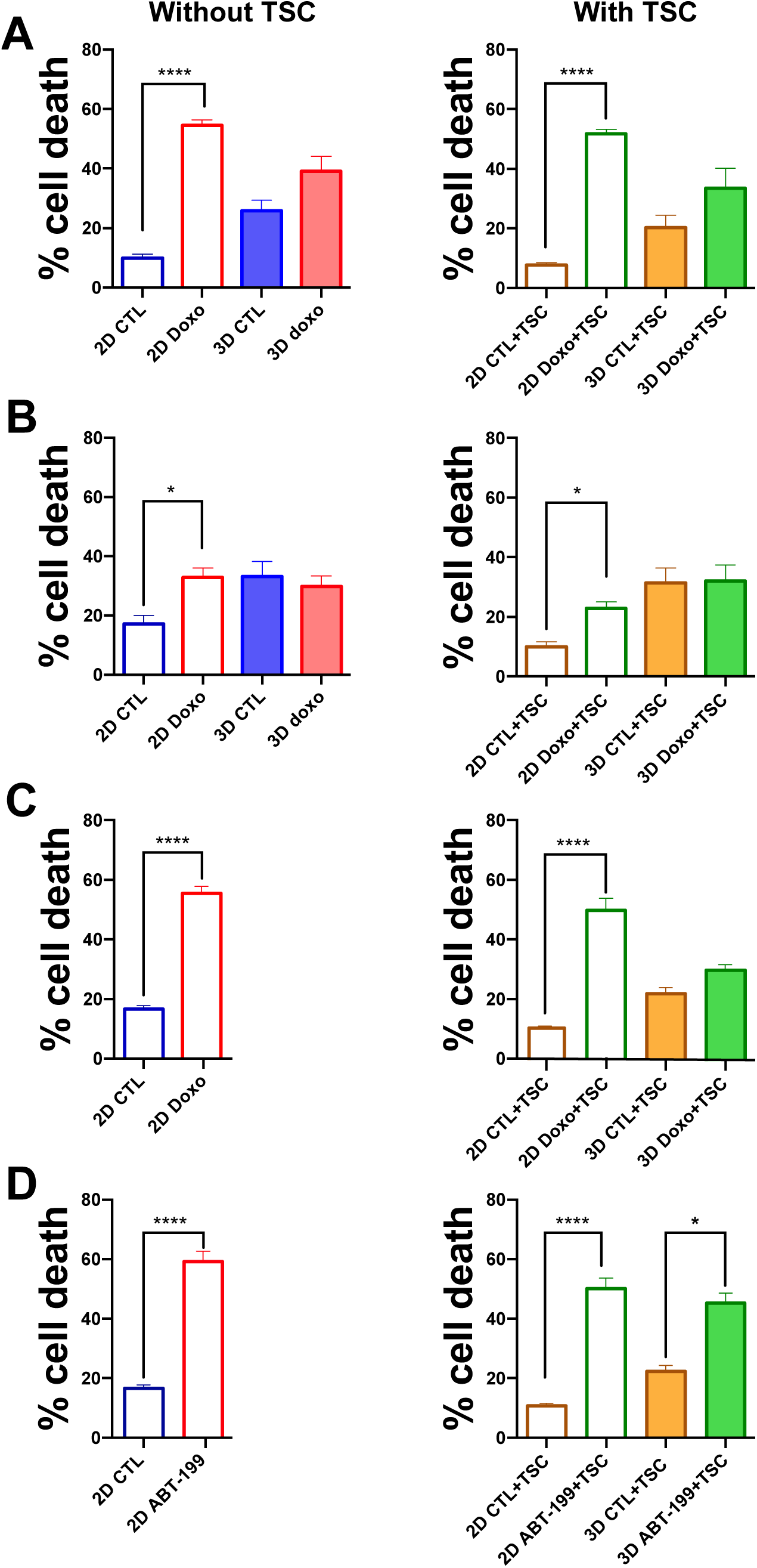
Impact of 3D culture on drug response in B-NHL. **A-C:** Comparison of cell death induced by doxorubicin in SU-DHL-4 (**A**), HLY1 (**B**) and DOHH2 (**C**) cultured in 2D or in 3D alone *(left panel)* or with TSC (2D+TSC or 3D+TSC) *(right panel).* For 2D culture, B cells were seeded alone or in presence of TSC. For 3D culture, 100 confluent capsules, containing B cells alone or B cells with TSC in presence of Matrigel, were seeded. 2D and 3D cultures were exposed to doxorubicin at a concentration corresponding to its EC50and cell death was evaluated after 24h.. **D:** Comparison of cell death induced by ABT-199 (1μM) in DOHH2 cells cultured in 2D *(left panel)* or in 3D with TSC and Matrigel *(right panel),* for 24h. The data represent mean ± SE of, at least, three independent experiments (Wilcoxon test. *P<0.05, **P<0.01, ***P<0.001, ****P<0.0001).

Altogether, these results reveal that our 3D lymphoma model may be useful to perform precision medicine or drug screening.

## DISCUSSION

Lymphomas develop as complex cell structures including tumor cells and their microenvironment within mechanically constraint lymph nodes (LN). Previous lymphoma spheroid models incorporating only malignant B cells (Decaup et al., 2013; Gravelle et al., 2014) have already suggested that the 3D cell architecture is a key feature for the regulation of lymphoma growth and therapeutic response. LN are highly dynamic structures expanding and becoming mechanically stiff under immune cell recruitment and proliferation, whereas immune response resolution is associated with LN contraction and return to a baseline of mechanical softness (Fletcher et al., 2015). LN mechanical properties rely on several determinants including the external constraint of the capsule and the internal tension created by the TSC network of specialized myofibroblasts able to produce and contract ECM. Moreover, mechanosensing has been recently demonstrated to control T-cell activation and metabolism (Meng et al., 2020) together with mechanical forces are involved in TCR (Hu and Butte, 2016) and BCR (Apoorva et al., 2018) signaling. Mouse models of FL and DLBCL cannot entirely recapitulate the biology of these tumors (Ramezani-Rad and Rickert, 2017) and, due to their inherent complexity and cost, they are not suitable to study specific cellular interactions contributing to the organization of lymphoma cell niche or to perform drug screening. The development of 3D lymphoma models including mechanical constraints is therefore of particular relevance to advance the understanding and treatment of these neoplasia. In the present work, we adapted a microfluidic approach to recapitulate the lymphoma microenvironment in hollow, permeable alginate shells, forming high-throughput (∼5000/s), size-controlled, and easy to handle spheroids (Alessandri et al., 2013). Such a strategy is ideally suited for drug testing and the versatility of the technique allows generating 3D monocultures (i.e. from tumor B cells only) or co-cultures with human stromal cells, and to evaluate the impact of ECM.

First, the proliferation of malignant B cells alone within spheroids results in cohesive multicellular aggregates promoted by the over-secretion of ECM in 3D compared to 2D cultures. Although expression and secretion of ECM have been mainly described in stromal cells, we demonstrated here that lymphoma B cells display a similar ability, as already proposed for normal plasmablasts (Della-Torre et al., 2020) and plasma cells (Tancred et al., 2009), or leukemic B cells (Mikaelsson et al., 2005). How ECM expression is regulated remains unclear. We could however hypothesize that it may originate from the activation of a mechanotransduction pathway as already described in other cell types (Coplen et al., 2003; He et al., 2004). Remarkably, alginate capsules are elastic, meaning that cell growth past confluence dilates the capsule, which conversely applies a compressive force onto the spheroid (Alessandri et al., 2013). Although not investigated in depth, we qualitatively observed that the cohesion of spheroids was stronger for post-confluent capsules, which would support this mechanotransduction hypothesis.

Second, as previously reported using other 3D models, some B-cell lines, including DOHH2, were unable to grow and form spheroids when maintained alone in capsules, reproducing the strong dependency of FL on their niche for survival and growth. Guided by our previous studies showing that lymphoid-like stromal cells have a supportive effect on FL B cells (Ame-Thomas et al., 2007; Gallouet et al., 2014; Guilloton et al., 2012), we added TSC to generate hybrid stroma-tumor 3D lymphoma models. Interestingly, our 3D model reproduced the different levels of NHL microenvironment dependency, mimicking the biology of aggressive versus indolent lymphomas (Scott and Gascoyne, 2014). By contrast with other 3D culture approaches that consist in embedding cells in an ECM scaffold (Gravelle et al., 2014; Purwada et al., 2015; Sabhachandani et al., 2019; Tian et al., 2015), our alginate spheroid model allows cell motions and interactions, leading to an early self­organization of a stroma-tumor B cell network that could organize localized 3D niches promoting cell interactions within the spheroids. Besides favoring or allowing malignant B-cell growth, the 3D co-culture was also associated with coevolution of stromal cells. In particular, TSC displayed increased podoplanin expression in 3D, mimicking the phenotype of FL-infiltrating stromal cells (Pandey et al., 2017), whereas such induction was not observed in 2D co-culture. Podoplanin is a transmembrane glycoprotein known to mediate the lymphoid stromal cell contractile function within LN (Astarita et al., 2015). It is thus tempting to speculate that podoplanin upregulation could contribute to the regulation of spheroid mechanical constraints. Production of TNF by malignant B cells has been proposed as a key mechanism of their dynamic interaction with stromal cells, leading to the upregulation of podoplanin, CCL2, and IL-8 in 2D culture conditions *in vitro* (Guilloton et al., 2012). However, neither TNF nor LT were found upregulated in DOHH2 maintained in 3D culture (data not shown), confirming a recent work performed on composite hydrogel-based lymphoma spheroids (Sabhachandani et al., 2019), suggesting that other soluble factors, but also 3D architecture and spatial constraints could contribute to the reprogramming of stromal cells.

Overall, the analysis of B-cell gene expression profile within spheroids revealed a profound shift associated with the transition from a 2D to a 3D context, mimicking the phenotype of primary FL B cells. The drop of B-cell proliferation that took place when cells filled the capsule was associated with an upregulation of quiescence genes also strongly expressed by FL B cells in agreement with their indolent phenotype. Even in the absence of a necrotic core within spheroids, the increase of *EGLN1* and *VEGF* expression could reflect a response to hypoxia, a process associated with a limitation of energy consumption through reduction of cell proliferation and DNA repair (Goda et al., 2003; Koshiji et al., 2004). Such association of response to hypoxia and quiescence has been previously described in a 3D model where spheroids were generated by the hanging drop method with lymphoma Β cells alone (Gravelle et al., 2014). Interestingly, BCL2 was found upregulated in 3D spheroids and has been previously shown to display anti­proliferative effects in normal and malignant Β cells (Tellier et al., 2014; Zinkel et al., 2006), suggesting that it could contribute to the decrease of cell proliferation. Even if the t(14;18) is the main driver of BCL2 deregulation in FL, BCL2 expression shows a high level of inter-and intra-tumoral heterogeneity and is correlated with IgH expression and NF-κΒ activation (Barreca et al., 2014; Heckman et al., 2002). Accordingly, we highlighted an upregulation of *RELB* and NF-κΒ target genes in 3D co-cultures, indicating that the NF-κΒ pathway was activated in lymphoma spheroids. Of note, a recent study revealed that fluid shear stress upregulates BCR expression and signaling in ABC-DLBCL (Apoorva et al., 2018), suggesting that engagement of BCR/NF-κB/BCL2 loop by biophysical forces could be a general feature of Β-cell lymphomas. Finally, we described in our 3D model an upregulation of *CXCR4,* previously associated with FL Β-cell migration, adhesion, and survival. Together with the reduction of *RGS13* involved in the organization of GC into dark and light zones and the limitation of GC size (Hwang et al., 2013; Shi et al., 2002), this could reproduce some of the factors sustaining FL dissemination and will require further investigations.

BCL2 expression is a key determinant of response to therapy in FL patients (Barreca et al., 2014). Moreover, the 3D-induced drug resistance has been already reported in solid cancers and hematological malignancies (Desoize, 2000; Gravelle et al., 2014; Tian et al., 2015). We confirmed here that the sensitivity to doxorubicin dramatically dropped when cells were cultured in 3D compared to 2D in all investigated B NHL cell lines. By contrast, the pro-apoptotic activity of ABT-199, mediating the dissociation of BCL2 with the proapoptotic BH3 only proteins, was maintained in 3D co-cultures despite BCL2 upregulation suggesting that ABT-199 might be considered as a good alternative for FL patient treatment. More generally, the cellular capsule technology adapted for lymphoma 3D culture seems a useful tool to screen new therapeutic approaches.

In summary, we report the development of a new lymphoma 3D model, including stromal cells and ECM, and recapitulating the dynamic crosstalk between tumor cells and their supportive microenvironment. The versatility of this high-throughput platform makes it possible to evaluate the role of various ECM components, stromal cell subtypes, and soluble factors on the spatial organization and function of the lymphoma niche. Moreover, this approach is particularly suitable for the testing of new therapeutic agents in B-NHL.

## MATERIALS AND METHODS

### Reagents and antibodies

Alginic acid sodium salt from brown algae, D-Sorbitol and CaCh hexahydrate, EGTA, paraformaldehyde were supplied by Merck (Fontenay sous bois, France). Matrigel was provided by Corning (UGAP, France). Propidium iodide and Phalloidin-Alexa 594 were from Thermo Fisher Scientific (Courtaboeuf, France). Hoechst 33258 and ABT-199 were purchased from Tocris Bioscience (Lille, France). The comprehensive cancer center pharmacy department of the Institut Bergonié kindly provided doxorubicin. APC Annexin V was from BD Biosciences (Le pont de Claix, France).

Rabbit polyclonal anti-laminin, rabbit polyclonal anti-fibronectin, rabbit monoclonal anti-human collagen I and chicken polyclonal anti-GFP antibodies were supplied by Abcam (Paris, France). Mouse anti-human CD20 APC-H7 and mouse anti-human CD73 PE were from e-Bioscience (Thermo Fisher Scientific). Monoclonal mouse anti-human Ki67 was from Dako Agilent Technologies (Les Ulis, France) and rabbit polyclonal anti-cleaved caspase 3 antibody was provided by Cell Signaling Technology (Ozyme, Saint Cyr l’école, France). All Alexa-conjugated secondary antibodies were purchased from Thermo Fisher Scientific.

### Encapsulation procedure

Cell encapsulation was performed by adapting the protocols from (Alessandri et al., 2016; Doméjean et al., 2016; Trushko et al., 2020). Briefly, it includes the fabrication of the device, the preparation of the solutions and the operation itself for cellular capsules formation.

First, the microfluidic device was printed with EnvisionTEC Micro Hi-Res Plus 3D printer (Arketyp3D, Le Raincy, France) using the resin HTM140V2 (EnvisionTEC), with the following printing parameters: burn-in range thickness 400μm, base plate of 300μm, and exposure time 3000ms. The printed device was washed using propanol and air-dried. In parallel, a glass capillary was pulled using a micropipette puller (P30, Sutter Instrument, Micro Mecanique SAS, Saint Germain en Laye, France), then cut and polished with micro-abrasive films (1μm grain, 3M) to reach the desired tip diameter (typically between 130 and 180 μm). This glass tip, which serves as a nozzle for the encapsulation device, was hydrophobized (1H,1H,2H,2H-Perfluorooctyltrimethoxysillane, ABCR) following standard protocols (Perret et al., 2002) and glued to the exit of the microfluidic plastic device with epoxyglue EA M-31CL (Loctite).

To run an encapsulation, the microfluidic device is connected to three syringes (10MDR-LL-GT SGE, Analytical Science, Milton Keynes, UK) mounted on pumps (Nemesys, Cetoni GmbH, Korbussen, Germany) for flow rate control. Three different solutions are injected into the three coaxial cones of the device. The outermost cone contains the alginate solution, which is prepared by dissolving 2% wt/vol sodium alginate (Protanal LF200S) in water and by adding 0.5mM SDS surfactant (VWR International, Fontenay sous bois, France). The solution was filtered at 1μm (glass filters, Pall Life Science) and stored at 4 °C. The intermediate cone contains a 300 mM sorbitol (Merck) solution. The innermost cone contains cells suspended in a sorbitol solution or cells/Matrigel/sorbitol solution (when ECM is needed) in a ratio of 2:1:2 (v/v). Lymphoma B cells lines were taken from cell suspension routinely cultured while adherent stromal cells were detached from the culture flask with trypsin-EDTA (Thermo Fisher Scientific). After rinse, cells were centrifuged and counted. For encapsulation of lymphoma cells or stromal cells only (mono-culture), a suspension of 2χ10^6^ cells were resuspended in culture DMEM (Thermo Fisher Scientific). When adherent and lymphoma Β cells were co-encapsulated, a suspension of 2*10^6^ cells of each cell type were used. 200μL of the cell suspension is injected into a cooling part to maintain Matrigel liquid (Alessandri et al., 2016). The flow rates are set to 45mL/h, 45mL/h and 30mL/h for the alginate, sorbitol and cell suspension, respectively. The device is positioned 40-50 cm above a petri dish containing 100mM CaCl_2_ solution for capsule gelling and collection. To improve capsule shape and monodispersity an alginate charging part and copper ring (21mm OD, RadioSpares), both connected to a high voltage (2 kV) generator (Stanford Research PS350, Acal BFI, France) are introduced, as described in (Trushko et al., 2020). The charged formed droplets passing through the copper ring under electrical tension get deflected as they cross the ring, creating a splay-like jet that prevents capsule coalescence. Once formed inside the calcium bath, capsules are washed with DMEM and transferred to cell culture medium.

### Cell lines, suspension, monolayer and encapsulated cell culture

Stromal cells (TSC) were isolated as previously described (Ame-Thomas et al., 2007; Guilloton et al., 2012) from human tonsils collected from children undergoing routine tonsillectomy, considered as chirurgical wastes. GFP-expressing TSC cells were established after transduction with lentivirus carrying the pUltra (gift from Malcolm Moore (Addgene plasmid # 24129; http://n2t.net/addgene:24129; RRID: Addgene_24129), which contained the enhanced GFP gene under the control of Ubc promoter and produced as previously described (WU and LU, 2007). TSC were transduced in RPMI supplemented with 2% FBS containing pUltra concentrated lentiviral supernatant with MOI of 4. On the next day, the medium was replaced with standard culture medium. After one week, eGFP^high^ TSC were isolated using a cell sorter (FACS Aria, BD Biosciences). For all the experiments reported in the present work, TSC were used from P9 to P13. DLBCL SU-DHL-4 and FL DOHH2 cell lines were obtained from the DSMZ cell collection (Braunschweig, Germany). HLY-1 cell line was kindly provided by Dr F. Meggetto (CRCT, Toulouse, France). Cells were routinely cultured in RPMI medium (Thermo Fisher Scientific), supplemented with 10% FBS (Dutscher, Brumath, France) under humid atmosphere containing 5% CO_2_, with medium changed 3 times a week. One day prior to encapsulation, cells were cultured in DMEM supplemented with 10% FBS. Once cellular capsules were formed following the protocol described above, they were cultured in 75 cm^2^ flasks in DMEM supplemented with 10% FBS under humid atmosphere containing 5% CO_2_, with medium changed 2 times a week. Depending on the type of experiments, capsules were sampled and transferred to dedicated culture supports, e.g. for imaging, or treated in bulk conditions and subsequently analyzed, e.g. for FACS.

### Spheroid growth measurement

At various post-encapsulation culture times, we selected about 200 capsules. Capsules were dissolved by incubation in PBS containing 1 mM EGTA and the spheroids were mechanically dissociated by pipetting. When spheroids were too cohesive, trypsin-EDTA was added to improve cell dissociation. The cell suspension was centrifuged and re-suspended in 200 μL of PBS and 50 μL of Precision Count Beads (BioLegend, Ozyme). The number of cells was then determined in a given cell suspension volume of 100 μL using flow cytometry (FACS Calibur, BD biosciences) according to the manufacturer’s instructions. To evaluate the cell death in spheroids, propidium iodide (PI) (10 μg/mL) was added to the cell suspension and PI fluorescence was analyzed by flow cytometry using Cell Quest or FlowJo (BD biosciences).

### Immunofluorescence staining of spheroids

#### In capsule staining

Capsules were fixed in DMEM w/o phenol red (Pan Biotech, Dutscher), containing 4% paraformaldehyde for 20 min at room temperature (RT). After rinsing, capsules were incubated for 30 min in DMEM w/o phenol red/ 0.1% triton X-100. Following a step of saturation with a solution containing DMEM w/o phenol red/ BSA 1% /FBS 2% for 1h at RT, capsules were incubated with primary antibody overnight at 4°C under shaking. Primary antibody was revealed by incubation of capsules with secondary Alexa-coupled antibody for 2h at RT. Nuclei were stained by incubation of capsules with Hoechst 33258 for 30 min at RT. To determine the role of Matrigel in the stromal network establishment, immunofluorescence using a rabbit anti-human pan laminin antibody revealed by goat anti rabbit Alexa 633 secondary antibody, was performed to visualize Matrigel, stromal cells were GFP positive and nuclei was stained with Hoechst 33258. Z-stacks were acquired using a Zeiss LSM 510 meta confocal microscope with a 25x objective (N.A 0.7). 3D rendering was obtained using UCSF Chimera software (RBVI, San Francisco, USA).

#### FFPE staining

To analyze protein expression inside spheroids, immunofluorescence on FFPE (Formalin-Fixed Paraffin-Embedded) spheroids sections (3-5 μm thick) were performed. Alginate shells were dissolved and spheroids were fixed as described above. Spheroids were then pre-embedded in 2% agarose and further embedded in a paraffin block. Immunofluorescence staining was then performed as previously described (Andrique et al., 2019). Images were acquired using a Zeiss LSM 510 meta confocal microscope (Zeiss, Göttingen, Germany) with a 25x objective (N.A 0.7). Then images obtained were processed using Fiji.

### 3D cell morphology quantification

Capsules were fixed at various post-encapsulation culture times and each capsule was placed in an agarose well of the Universlide microscopy chamber (Alessandri et al., 2017), which allowed immobilization of the capsules as required for stable z-stack acquisition and medium throughput 3D imaging. Cell nuclei were stained with Hoechst 33258 and stromal cells were visualized by the GFP expression. Images were acquired using a Leica SP8 confocal microsocpe with a 20X dry objective (NA 0.7). To measure the evolution of TSC complexity over time two parameters were measured: the sphericity defined as 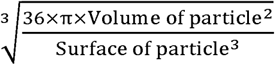 and the branchness defined as 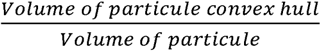. To obtain these values, we applied a locally adjusted threshold to the green signal, performed 3D segmentation (3D suite plugin, Fiji (Ollion et al., 2013). For the two parameters, the values of all “particles” contained in one capsule were averaged. Thus, each capsule was characterized by one mean value for sphericity and branchness. 10-18 capsules were analyzed per time point

### Multiplex fluorescence microscopy

Patients were recruited under institutional review board approval following informed consent process according to the declaration of Helsinki and the French National Cancer Institute ethic committee recommendations. Tissue samples included tonsils collected from routine tonsillectomy, and lymph nodes obtained from grade 1-2 FL and from DLBCL patients.

Four-micrometer-thick whole-slide sections, obtained with a microtome (Histocore multicut Leicabiosystems, Nanterre, France) from FFPE tissue or spheroids blocks, were transferred onto plus-charged slides (VWR international), followed by multiplex immunofluorescence staining with a U DISCOVERY 5 plex immunofluorescence (Roche Diagnostics, Meylan, France). Three sequential rounds of staining were performed each including heat deactivation step, followed by incubation with primary antibody and corresponding HRP secondary antibody. Hence, rat anti-podoplanin (Thermo Fisher Scientific NZ-1.3), mouse Anti-PAX5 antibody (Abcam ab224660), and sheep anti-transglutaminase (R&D AF4376) expressions were visualized on the same section using HRP enzyme mediated deposition of the tyramide coupled to rhodamine, DCC and FAM fluorophores (kits Ventana Medical Systems, Tucson, Ariz), respectively, that covalently binds to the tissue at the site of the reaction. After 3 sequential reactions, sections were counterstained with DAPI and coverslipped using fluoromount (Enzo Life Sciences, Farmingdale, NY, USA). Visualization was performed with the Nanozoomer (Hamamatsu Photonics, Massy, France) equipped with the multicolor fluorescence module.

### Cell co-culture and cell sorting

2D co-cultures were performed over 3 days. 3*10^5^DOHH2 (1.5*10^5^/mL) were plated on 5*10^4^ TSC /cm^2^ in 6 well plates. For cell sorting, cells were trypsinized (Thermo Fisher Scientific) for 12 min at 37 °C and washed with medium DMEM 10% FBS. 3D co-cultures were performed over 10 days. For cell sorting, after dissolution of the alginate shells, cells were rinsed with DMEM 10% FBS and centrifuged for 7 min at 1700 rpm, incubated with trypsin at 37 °C for 15 min, and rinsed with medium DMEM 10% FBS. Cells were stained for 20 min on ice with mouse anti-human CD20 APC-H7 and mouse anti-human CD73 PE. After DAPI staining, cell sorting was processed by the FACSAria II (BD Biosciences).

### RT q-PCR arrays

After cell sorting, RNA extraction from DOHH2 co-cultured in 2D or 3D with TSC cells was done using Nucleospin RNA XS Micro kit (PN 740990) from Macherey Nagel (Hoerdt, France). cDNA were synthesized using reverse transcription master mix (Fluidigm, USA, PN 100-6297), followed by gene expression preamplification with Fluidigm Preamp Master mix (PN 100-5744) and Taq Man assays and followed by RT-QPCR. Assay-on-demand primers, probes, and TaqMan gene expression Master Mix were obtained from Thermo Fisher Scientific. Gene expression was measured using 96.96 Dynamic Array^™^ IFCs and the BioMark^™^ HD System from Fluidigm, based on ΔCt calculation method. A tonsil pool was used as appropriate DOHH2 expression external control. For each sample, the Ct value for the gene of interest was determined and normalized to its respective mean value of housekeeping genes *(ABL1* and *GAPDH).* Data were represented through a heatmap using the pheatmap R package. In addition, differentially expressed genes were analyzed in Affymetrix data sets comparing purified FL Β cells and purified centrocytes (GSE85233 and GSE136249) (Pangault et al., 2020).

### Chemotherapeutic toxicity measurement

To compare the drug toxicities between lymphoma cells cultured in suspension and in spheroids, 0.5×10^6^cells/well co-cultured or not with 5χ10^4^ TSC /cm^2^ or 100 confluent spheroids were distributed in 6-well plates and treated or not with doxorubicin for 24h. Then, cell death was assessed by flow cytometry after Annexin V-APC labelling according to the manufacturer’s instructions.

### Statistical analyses

All data were expressed as means ± SEM. The significance of differences was calculated using the parametric ANOVA or the non-parametric U-Mann and Whitney or Wilcoxon tests, as appropriate. (*P<0.05, **P<0.01, ***P<0.001, ****P<0.0001). Statistical analyses were performed using GraphPad Prism 8 software.

## ACKNOWLEDGMENTS

We thank Stephanie Durand-Panteix for her technical support with transduction experiments. We thank Atika Zouine and Vincent Pitard for technical assistance at the Flow cytometry facility, CNRS UMS 3427, INSERM US 005, Univ. Bordeaux, F-33000 Bordeaux, France. We acknowledge the Bordeaux Imaging Center, a service unit of the CNRS-INSERM and Bordeaux University, member of the national infrastructure France BioImaging supported by the French Research Agency (ANR-10-INBS-04). A part of immunofluorescence study was performed on the Microscopy Rennes Imaging Center (MRic-ALMF; UMS 6480 Biosit, Rennes, France) member of the national infrastructure France-BioImaging. Cell sorting was performed at the Biosit Flow Cytometry and Cell Sorting Facility CytomeTRI (UMS 6480 Biosit, Rennes, France).

---

**Movie 1: Dissolution of alginate shell.** Alginate shell of capsule containing SU-DHL-4 spheroid obtained after 10 days in culture is dissolved in a bath of PBS/EGTA. The images were acquired every 15 seconds until complete dissolution of capsule using an epifluorescence microscope (Leica DMI8).

**Movie 2: Establishment of the stromal cell network.** TSC expressing GFP and DOHH2 expressing dTomato were co-encapsulated in presence of Matrigel and placed in an Universlide in DMEM containing 10% FCS. The images were acquired immediately after encapsulation every 30 minutes for 12 hours using an epifluorescence microscope (Leica DMI8) equipped with a thermostatic and CO2 chamber (Live on Stage, Leica). Scale bar: 100μm.

**Movie 3: Spatial organization of lymphoma spheroid.** 3D reconstruction from optical sections using confocal microscopy of capsule containing DOHH2 with Mg and TSC cultured for 6 days.

## REFERENCES

Alessandri, K., Andrique, L., Feyeux, M., Bikfalvi, A., Nassoy, P., Recher, G., 2017. All-in-one 3D printed microscopy chamber for multidimensional imaging, the UniverSlide. Sci.Rep. 7, 42378. https://doi.org/10.1038/srep42378

Alessandri, K., Feyeux, M., Gurchenkov, Β., Delgado, C., Trushko, A., Krause, K.H., Vignjevic, D., Nassoy, P., Roux, A., 2016. A 3D printed microfluidic device for production of functionalized hydrogel microcapsules for culture and differentiation of human Neuronal Stem Cells (hNSC). Lab Chip. 16, 1593–1604. https://doi.org/10.1039/c6lc00133e

Alessandri, K., Sarangi, B.R., Gurchenkov, V.V., Sinha, Β., Kiessling, T.R., Fetler, L., Rico, F., Scheuring, S., Lamaze, C., Simon, A., Geraldo, S., Vignjevic, D., Domejean, H., Rolland, L., Funfak, A., Bibette, J., Bremond, N., Nassoy, P., 2013. Cellular capsules as a tool for multicellular spheroid production and for investigating the mechanics of tumor progression in vitro. P roc. Natl. Acad. Sci. U.S. A 110, 14843–14848. https://doi.org/10.1073/pnas.1309482110

Alizadeh, A.A., Eisen, M.B., Davis, R.E., Ma, C., Lossos, I.S., Rosenwald, A., Boldrick, J.C., Sabet, H., Tran, T., Yu, X., Powell, J.I., Yang, L., Marti, G.E., Moore, T., Hudson, J., Lu, L., Lewis, D.B., Tibshirani, R., Sherlock, G., Chan, W.C., Greiner, T.C., Weisenburger, D.D., Armitage, J.O., Warnke, R., Levy, R., Wilson, W., Grever, M.R., Byrd, J.C., Botstein, D., Brown, P.O., Staudt, L.M., 2000. Distinct types of diffuse large Β-cell lymphoma identified by gene expression profiling. Nature 403, 503–511. https://doi.org/10.1038/35000501

Ame-Thomas, P., Maby-El, H.H., Monvoisin, C., Jean, R., Monnier, D., Caulet-Maugendre, S., Guillaudeux, T., Lamy, T., Fest, T., Tarte, K., 2007. Human mesenchymal stem cells isolated from bone marrow and lymphoid organs support tumor B-cell growth: role of stromal cells in follicular lymphoma pathogenesis. Blood 109, 693–702. https://doi.org/10.1182/blood-2006-05-020800

Andrique, L., Recher, G., Alessandri, K., Pujol, N., Feyeux, M., Bon, P., Cognet, L., Nassoy, P., Bikfalvi, A., 2019. A model of guided cell self-organization for rapid and spontaneous formation of functional vessels. Science Advances 5. https://doi.org/10.1126/sciadv.aau6562

Apoorva, F., Loiben, A.M., Shah, S.B., Purwada, A., Fontan, L., Goldstein, R., Kirby, B.J., Melnick, A.M., Cosgrove, B.D., Singh, A., 2018. How Biophysical Forces Regulate Human B Cell Lymphomas. Cell Reports 23, 499–511. https://doi.org/10.1016/j.celrep.2018.03.069

Astarita, J.L., Cremasco, V., Fu, J., Darnell, M.C., Peck, J.R., Nieves-Bonilla, J.M., Song, K., Kondo, Y., Woodruff, M.C., Gogineni, A., Onder, L., Ludewig, B., Weimer, R.M., Carroll, M.C., Mooney, D.J., Xia, L., Turley, S.J., 2015. The CLEC-2-podoplanin axis controls the contractility of fibroblastic reticular cells and lymph node microarchitecture. Nature Immunology 16, 75–84. https://doi.org/10.1038/ni.3035

Barreca, A., Martinengo, C., Annaratone, L., Righi, L., Chiappella, A., Ladetto, M., Demurtas, A., Chiusa, L., Stacchini, A., Crosetto, N., van Oudenaarden, A., Chiarle, R., 2014. Inter-and intratumoral heterogeneity of BCL2 correlates with IgH expression and prognosis in follicular lymphoma. Blood Cancer Journal 4, e249–e249. https://doi.org/10.1038/bcj.2014.67

Benien, P., Swami, A., 2014. 3D tumor models: history, advances and future perspectives. Future Oncol 10, 1311–1327. https://doi.org/10.2217/fon.13.274

Bissell, M.J., Radisky, D., 2001. Putting tumours in context. Nat.Rev.Cancer 1, 46–54. https://doi.org/10.1038/35094059

Chen, J., Alexander, J.S., Orr, A.W., 2012. Integrins and Their Extracellular Matrix Ligands in Lymphangiogenesis and Lymph Node Metastasis. Int J Cell Biol 2012. https://doi.org/10.1155/2012/853703

Coplen, D.E., Macarak, E.J., Howard, P.S., 2003. Matrix synthesis by bladder smooth muscle cells is modulated by stretch frequency. In Vitro Cell.Dev.Biol.-Animal 39, 157–162. https://doi.org/10.1007/s11626-003-0010-3

Daster, S., Amatruda, N., Calabrese, D., Ivanek, R., Turrini, E., Droeser, R.A., Zajac, P., Fimognari, C., Spagnoli, G.C., Iezzi, G., Mele, V., Muraro, M.G., 2016. Induction of hypoxia and necrosis in multicellular tumor spheroids is associated with resistance to chemotherapy treatment. Oncotarget 8, 1725–1736. https://doi.org/10.18632/oncotarget.13857

Decaup, E., Jean, C., Laurent, C., Gravelle, P., Fruchon, S., Capilla, F., Marrot, A., Al Saati, T., Frenois, F.-X., Laurent, G., Klein, C., Varoqueaux, N., Savina, A., Fournié, J.-J., Bezombes, C., 2013. Anti-tumor activity of obinutuzumab and rituximab in a follicular lymphoma 3D model. Blood Cancer J 3, e131. https://doi.org/10.1038/bcj.2013.32

Decaup, E., Rossi, C., Gravelle, P., Laurent, C., Bordenave, J., Tosolini, M., Tourette, A., Perrial, E., Dumontet, C., Poupot, M., Klein, C., Savina, A., Fournié, J.-J., Bezombes, C., 2019. A Tridimensional Model for NK Cell-Mediated ADCC of Follicular Lymphoma. Front. Immunol. 10. https://doi.org/10.3389/fimmu.2019.01943

Della-Torre, E., Rigamonti, E., Perugino, C., Baghai-Sain, S., Sun, N., Kaneko, N., Maehara, T., Rovati, L., Ponzoni, M., Milani, R., Lanzillotta, M., Mahajan, V., Mattoo, H., Molineris, I., Deshpande, V., Stone, J.H., Falconi, M., Manfredi, A.A., Pillai, S., 2020. B lymphocytes directly contribute to tissue fibrosis in patients with IgG4-related disease. Journal of Allergy and Clinical Immunology 145, 968–981.e14. https://doi.org/10.1016/j.jaci.2019.07.004

Desoize, B., 2000. Multicellular resistance: a paradigm for clinical resistance? Critical Reviews in Oncology/Hematology 36, 193–207. https://doi.org/10.1016/S1040-8428(00)00086-X

Domejean, H., Pierre, M. de la M.S., Funfak, A., Atrux-Tallau, N., Alessandri, K., Nassoy, P., Bibette, J., Bremond, N., 2016. Controlled production of sub-millimeter liquid core hydrogel capsules for parallelized 3D cell culture. Lab Chip 17, 110–119. https://doi.org/10.1039/C6LC00848H

Fletcher, A.L., Acton, S.E., Knoblich, K., 2015. Lymph node fibroblastic reticular cells in health and disease. Nat. Rev. Immunol. 15, 350–361. https://doi.org/10.1038/nri3846

Fowler, N.H., Cheah, C.Y., Gascoyne, R.D., Gribben, J., Neelapu, S.S., Ghia, P., Bollard, C., Ansell, S., Curran, M., Wilson, W.H., O’Brien, S., Grant, C., Little, R., Zenz, T., Nastoupil, L.J., Dunleavy, K., 2016. Role of the tumor microenvironment in mature B-cell lymphoid malignancies. Haematologica 101, 531–540. https://doi.org/10.3324/haematol.2015.139493

Gallouet, A.S., Travert, M., Bresson-Bepoldin, L., Guilloton, F., Pangault, C., Caulet-Maugendre, S., Lamy, T., Tarte, K., Guillaudeux, T., 2014. COX-2-independent effects of celecoxib sensitize lymphoma B cells to TRAIL-mediated apoptosis. Clin.Cancer Res. 20, 2663–2673. https://doi.org/10.1158/1078-0432.CCR-13-2305

Goda, N., Dozier, S.J., Johnson, R.S., 2003. HIF-1 in Cell Cycle Regulation, Apoptosis, and Tumor Progression. Antioxidants & Redox Signaling 5, 467–473. https://doi.org/10.1089/152308603768295212

Gravelle, P., Jean, C., Familiades, J., Decaup, E., Blanc, A., Bezombes-Cagnac, C., Laurent, C., Savina, A., Fournié, J.-J., Laurent, G., 2014. Cell growth in aggregates determines gene expression, proliferation, survival, chemoresistance, and sensitivity to immune effectors in follicular lymphoma. Am. J. Pathol. 184, 282–295. https://doi.org/10.1016/j.ajpath.2013.09.018

Guilloton, F., Caron, G., Menard, C., Pangault, C., Ame-Thomas, P., Dulong, J., De, V.J., Rossille, D., Henry, C., Lamy, T., Fouquet, O., Fest, T., Tarte, K., 2012. Mesenchymal stromal cells orchestrate follicular lymphoma cell niche through the CCL2-dependent recruitment and polarization of monocytes. Blood 119, 2556­2567. https://doi.org/10.1182/blood-2011-08-370908

He, Y., Macarak, E.J., Korostoff, J.M., Howard, P.S., 2004. Compression and Tension: Differential Effects on Matrix Accumulation by Periodontal Ligament Fibroblasts In Vitro. Connective Tissue Research 45, 28–39. https://doi.org/10.1080/03008200490278124

Heckman, C.A., Mehew, J.W., Boxer, L.M., 2002. NF-kappaB activates Bcl-2 expression int(14; 18) lymphoma cells. Oncogene 21, 3898–3908. https://doi.org/10.1038/sj.onc.1205483

Heldt, F.S., Barr, A.R., Cooper, S., Bakal, C., Novák, Β., 2018. A comprehensive model for the proliferation-quiescence decision in response to endogenous DNA damage in human cells. PNAS 115, 2532–2537.

Hu, K.H., Butte, M.J., 2016. T cell activation requires force generation. The Journal of cell biology 213, 535–542. https://doi.org/10.1083/jcb.201511053

Huet, S., Sujobert, P., Salles, G., 2018. From genetics to the clinic: a translational perspective on follicular lymphoma. Nat Rev Cancer 18, 224–239. https://doi.org/10.1038/nrc.2017.127

Hwang, I.-Y., Hwang, K.-S., Park, C., Harrison, K.A., Kehrl, J.H., 2013. Rgs13 Constrains Early B Cell Responses and Limits Germinal Center Sizes. PLoS ONE 8, e60139. https://doi.org/10.1371/journal.pone.0060139

Kennedy, R., Klein, U., 2018. Aberrant Activation of NF-**k**B Signalling in Aggressive Lymphoid Malignancies. Cells 7, 189. https://doi.org/10.3390/cells7110189

Koshiji, M., Kageyama, Y., Pete, E.A., Horikawa, I., Barrett, J.C., Huang, L.E., 2004. HIF-1α induces cell cycle arrest by functionally counteracting Myc. EMBO J 23, 1949–1956. https://doi.org/10.1038/sj.emboj.7600196

Meng, K.P., Majedi, F.S., Thauland, T.J., Butte, M.J., 2020. Mechanosensing through YAP controls T cell activation and metabolism. Journal of Experimental Medicine 217, e20200053. https://doi.org/10.1084/jem.20200053

Mikaelsson, E., Danesh-Manesh, A.H., Luppert, A., Jeddi-Tehrani, M., Rezvany, M.-R., Sharifian, R.A., Safaie, R., Roohi, A., Osterborg, A., Shokri, F., Mellstedt, H., Rabbani, H., 2005. Fibromodulin, an extracellular matrix protein: characterization of its unique gene and protein expression in B-cell chronic lymphocytic leukemia and mantle cell lymphoma. Blood 105, 4828–4835. https://doi.org/10.1182/blood-2004-10-3941

Ollion, J., Cochennec, J., Loll, F., Escudé, C., Boudier, T., 2013. TANGO: a generic tool for high-throughput 3D image analysis for studying nuclear organization. Bioinformatics 29, 1840–1841. https://doi.org/10.1093/bioinformatics/btt276

Pandey, S., Mourcin, F., Marchand, T., Nayar, S., Guirriec, M., Pangault, C., Monvoisin, C., Amé-Thomas, P., Guilloton, F., Dulong, J., Coles, M., Fest, T., Mottok, A., Barone, F., Tarte, K., 2017. IL-4/CXCL12 loop is a key regulator of lymphoid stroma function in follicular lymphoma. Blood 129, 2507–2518. https://doi.org/10.1182/blood-2016-08-737239

Pangault, C., Amé-Thomas, P., Rossille, D., Dulong, J., Caron, G., Nonn, C., Chatonnet, F., Desmots, F., Launay, V., Lamy, T., Fest, T., Tarte, K., 2020. Integrative Analysis of Cell Crosstalk within Follicular Lymphoma Cell Niche: Towards a Definition of the FL Supportive Synapse. Cancers 12, 2865. https://doi.org/10.3390/cancers12102865

Perret, E., Leung, A., Morel, A., Feracci, H., Nassoy, P., 2002. Versatile Decoration of Glass Surfaces To Probe Individual Protein-Protein Interactions and Cellular Adhesion. Langmuir 18, 846–854. https://doi.org/10.1021/la015601y

Purwada, A., Jaiswal, M.K., Ahn, H., Nojima, T., Kitamura, D., Gaharwar, A.K., Cerchietti, L., Singh, A., 2015. Ex vivo engineered immune organoids for controlled germinal center reactions. Biomaterials 63, 24–34. https://doi.org/10.1016/j.biomaterials.2015.06.002

Ramezani-Rad, P., Rickert, R.C., 2017. Murine models of germinal center derived-lymphomas. Current Opinion in Immunology, Lymphocyte development and activation * Tumour immunology 45, 31–36. https://doi.org/10.1016/j.coi.2016.12.002

Sabhachandani, P., Sarkar, S., Mckenney, S., Ravi, D., Evens, A.M., Konry, T., 2019. Microfluidic assembly of hydrogel-based immunogenic tumor spheroids for evaluation of anticancer therapies and biomarker release. Journal of Controlled Release 295, 21–30. https://doi.org/10.1016/jjconrel.2018.12.010

Scott, D.W., Gascoyne, R.D., 2014. The tumour microenvironment in Β cell lymphomas. Nature Reviews Cancer 14, 517–534. https://doi.org/10.1038/nrc3774

Sehn, L.H., Gascoyne, R.D., 2015. Diffuse large Β-cell lymphoma: optimizing outcome in the context of clinical and biologic heterogeneity. Blood 125, 22–32. https://doi.org/10.1182/blood-2014-05-577189

Sgambato, A., Cittadini, A., Faraglia, Β., Weinstein, I.B., 2000. Multiple functions of p27Kip1 and its alterations in tumor cells: a review. Journal of Cellular Physiology 183, 18–27. https://doi.org/10.1002/(SICI)1097-4652(200004)183:1<18::AID-JCP3>3.0.CO;2-S

Shi, G.-X., Harrison, K., Wilson, G.L., Moratz, C., Kehrl, J.H., 2002. RGS13 Regulates Germinal Center Β Lymphocytes Responsiveness to CXC Chemokine Ligand (CXCL)12 and CXCL13. J Immunol 169, 2507–2515. https://doi.org/10.4049/jimmunol.169.5.2507

Smith, A., Howell, D., Patmore, R., Jack, A., Roman, E., 2011. Incidence of haematological malignancy by sub-type: a report from the Haematological Malignancy Research Network. British Journal of Cancer 105, 1684–1692. https://doi.org/10.1038/bjc.2011.450

Sobocinski, G.P., Toy, K., Bobrowski, W.F., Shaw, S., Anderson, A.O., Kaldjian, E.P., 2010. Ultrastructural localization of extracellular matrix proteins of the lymph node cortex: evidence supporting the reticular network as a pathway for lymphocyte migration. BMC Immunol 11, 42. https://doi.org/10.1186/1471-2172-11-42

Sutherland, R.M., 1988. Cell and environment interactions in tumor microregions: the multicell spheroid model. Science 240, 177–184.

Tancred, T.M., Belch, A.R., Reiman, T., Pilarski, L.M., Kirshner, J., 2009. Altered Expression of Fibronectin and Collagens I and IV in Multiple Myeloma and Monoclonal Gammopathy of Undetermined Significance. J Histochem Cytochem 57, 239–247. https://doi.org/10.1369/jhc.2008.952200

Tellier, J., Menard, C., Roulland, S., Martin, N., Monvoisin, C., Chasson, L., Nadel, A., Gaulard, P., Schiff, C., Tarte, K., 2014. Human t(14;18)positive germinal center B cells: a new step in follicular lymphoma pathogenesis? Blood 123, 3462–3465. https://doi.org/10.1182/blood-2013-12-545954

Teras, L.R., DeSantis, C.E., Cerhan, J.R., Morton, L.M., Jemal, A., Flowers, C.R., 2016. 2016 US lymphoid malignancy statistics by World Health Organization subtypes. CA: A Cancer Journal for Clinicians 66, 443–459. https://doi.org/10.3322/caac.21357

Tian, Y.F., Ahn, H., Schneider, R.S., Yang, S.N., Roman-Gonzalez, L., Melnick, A.M., Cerchietti, L., Singh, A., 2015. Integrin-specific hydrogels as adaptable tumor organoids for malignant B and T cells. Biomaterials 73, 110–119. https://doi.org/10.1016/j.biomaterials.2015.09.007

Trushko, A., Di Meglio, I., Merzouki, A., Blanch-Mercader, C., Abuhattum, S., Guck, J., Alessandri, K., Nassoy, P., Kruse, K., Chopard, B., Roux, A., 2020. Buckling of an Epithelium Growing under Spherical Confinement. Developmental Cell 54, 655–668.e6. https://doi.org/10.1016/j.devcel.2020.07.019

Verdiere, L., Mourcin, F., Tarte, K., 2018. Microenvironment signaling driving lymphomagenesis. Current Opinion in Hematology 25, 335.

Wu, C., Lu, Y., 2007. Inclusion of high molecular weight dextran in calcium phosphate-mediated transfection significantly improves gene transfer efficiency. Cell Mol Biol (Noisy-le-grand) 53, 67–74.

Zhang, Jun, Seet, C.S., Sun, C., Li, J., You, D., Volk, A., Breslin, P., Li, X., Wei, W., Qian, Z., Zeleznik-Le, N.J., Zhang, Z., Zhang, Jiwang, 2013. p27kip1 maintains a subset of leukemia stem cells in the quiescent state in murine MLL-leukemia. Mol Oncol 7, 1069–1082. https://doi.org/10.1016/j.molonc.2013.07.011

Zinkel, S., Gross, A., Yang, E., 2006. BCL2 family in DNA damage and cell cycle control. Cell Death Differ. 13, 1351–1359. https://doi.org/10.1038/sj.cdd.4401987

