## Supplemental figures for "Stromal cells regulate malignant B-cell spatial organization, survival, and drug response in a new 3D model mimicking lymphoma tumor niche"

FIGURE S1

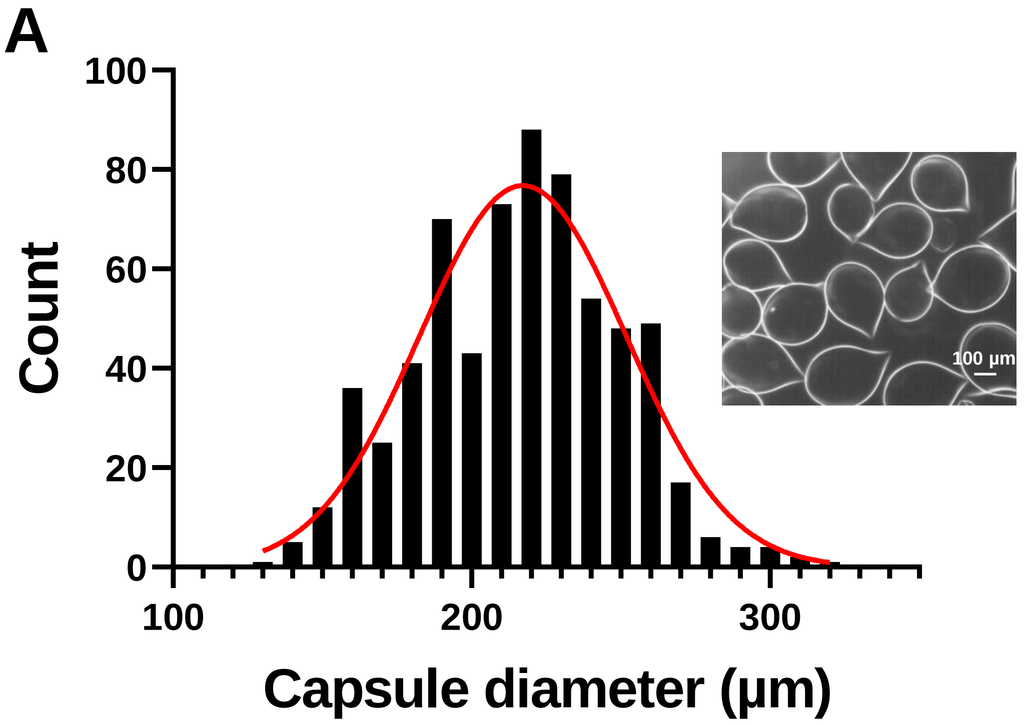


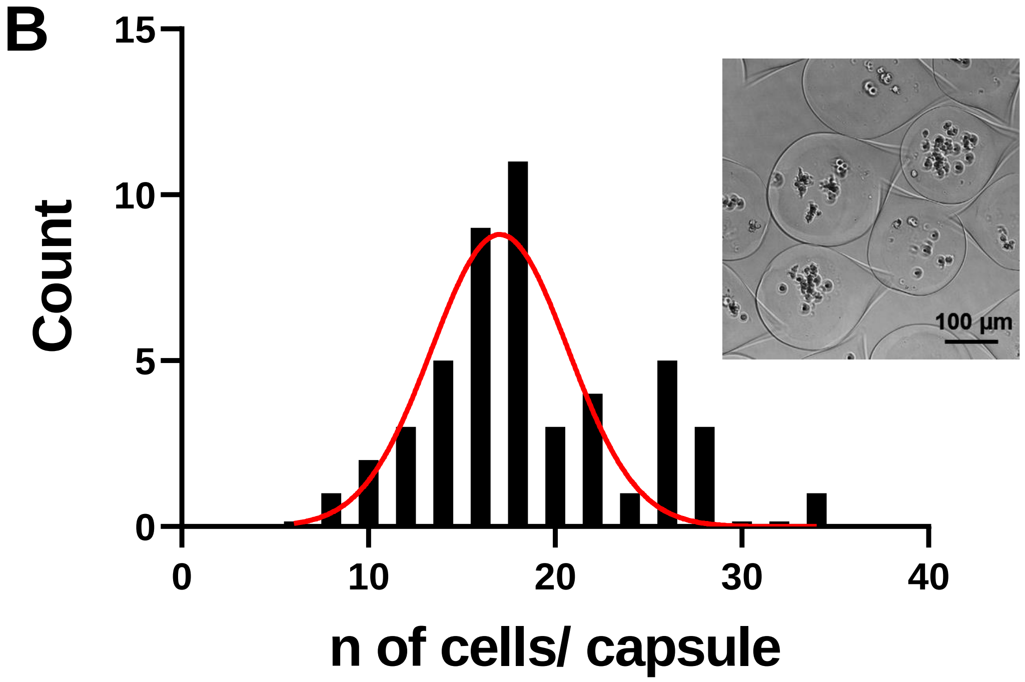


**Figure S1: Distribution of (A) the alginate microcapsule size and (B) the number of cells in capsules immediately after encapsulation.**

### FIGURE S2

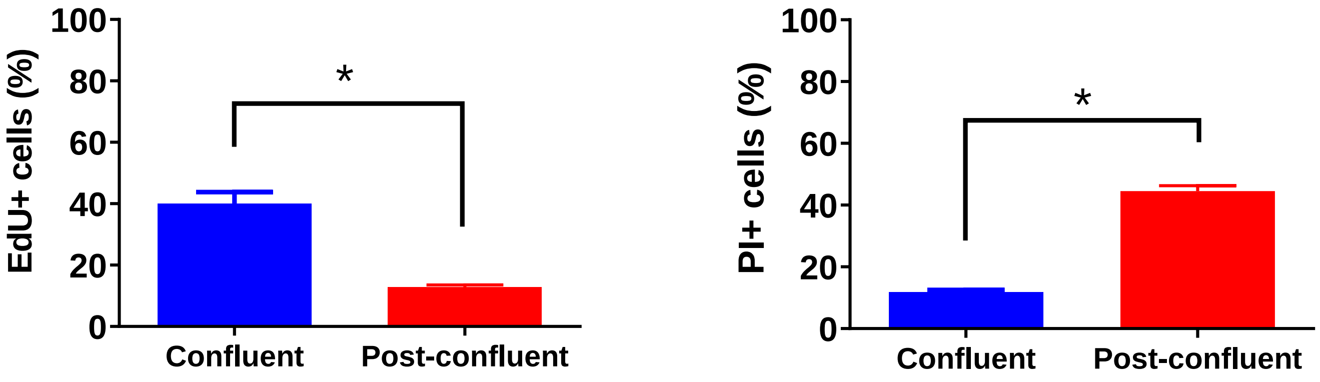


**Figure S2: Evaluation of cell proliferation and cell death in confluent and post-confluent capsules.** Proliferative cells were stained using Click-it EdU cell proliferation kit (ThermoFisher Scientific) as manufacturer's recommendations. Dead cells were detected by propidium iodide labelling. Then, the percentage of fluorescent proliferative (EdU+) or dead (PI+) cells were measured by flow cytometry.

### FIGURE S3

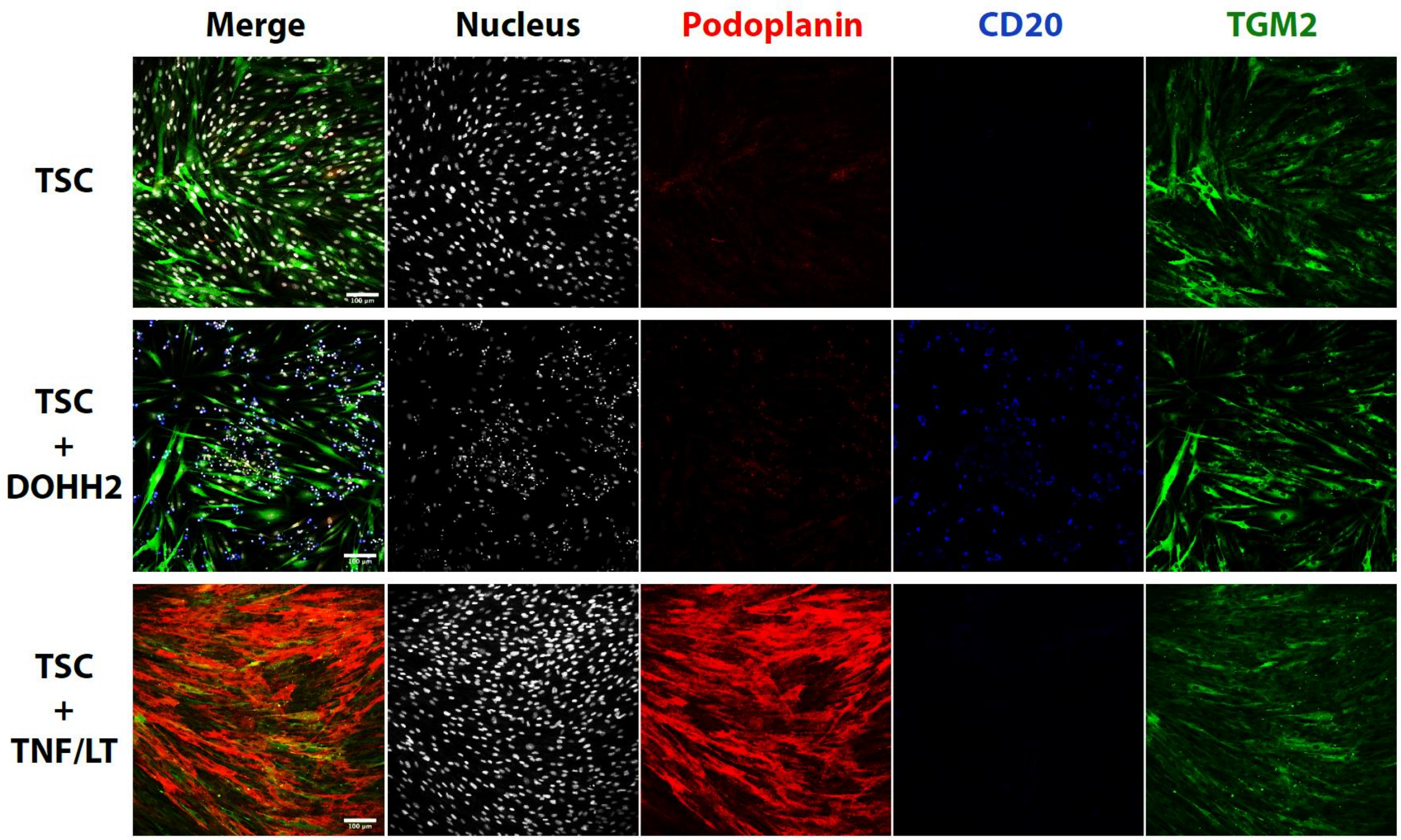


**Figure S3: Expression of lymphoid stromal markers by TSC cultured in 2D.**
TSCs were cultured with or without DOHH2 expressing CD20 (in blue). In these conditions TSCs express transglutaminase 2 (TGM2, in green) but not podoplanin (in red). Only the induction by TNF/LT lead to the expression of podoplanin (in red) by TSCs in 2D. Scale bar: 100µm

### FIGURE S4

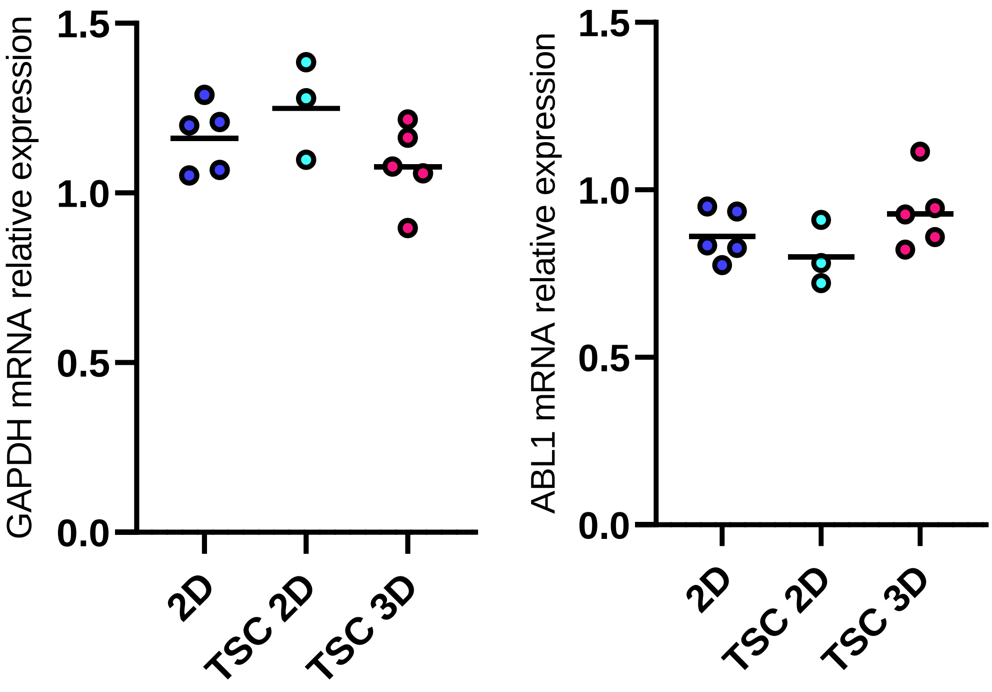


**Figure S4: Expression of GAPDH and ABL1 housekeeping genes** in DOHH2 cultured in 2D with (n=5) or without TSC (n=3) and in 3D with TSC (n=5) for 10 days. Results represent the median of independent experiments.

### FIGURE S5

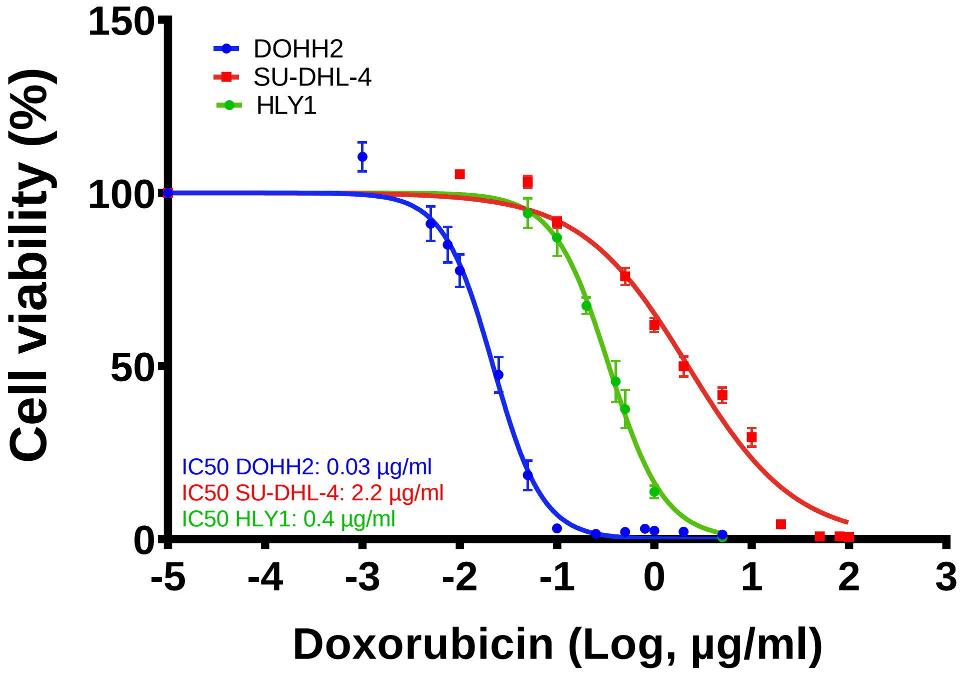


**Figure S5: Determination of the EC50 of various cell lines for doxorubicin.** Cell lines were plated in 96-well plates at 20,000 cells per well, with various concentrations of doxorubicin for 24h. The half maximal inhibitory concentration (IC50) was determined using the luminescent cell viability assay CellTiter-Glo® (Promega, USA). Luminescence levels were quantified using the FlexStation® 3 (Molecular Devices, USA).
