## Supplementary figures and images for "Stromal cells regulate malignant B-cell spatial organization, survival, and drug response in a new 3D model mimicking lymphoma tumor niche"

### Movie 1

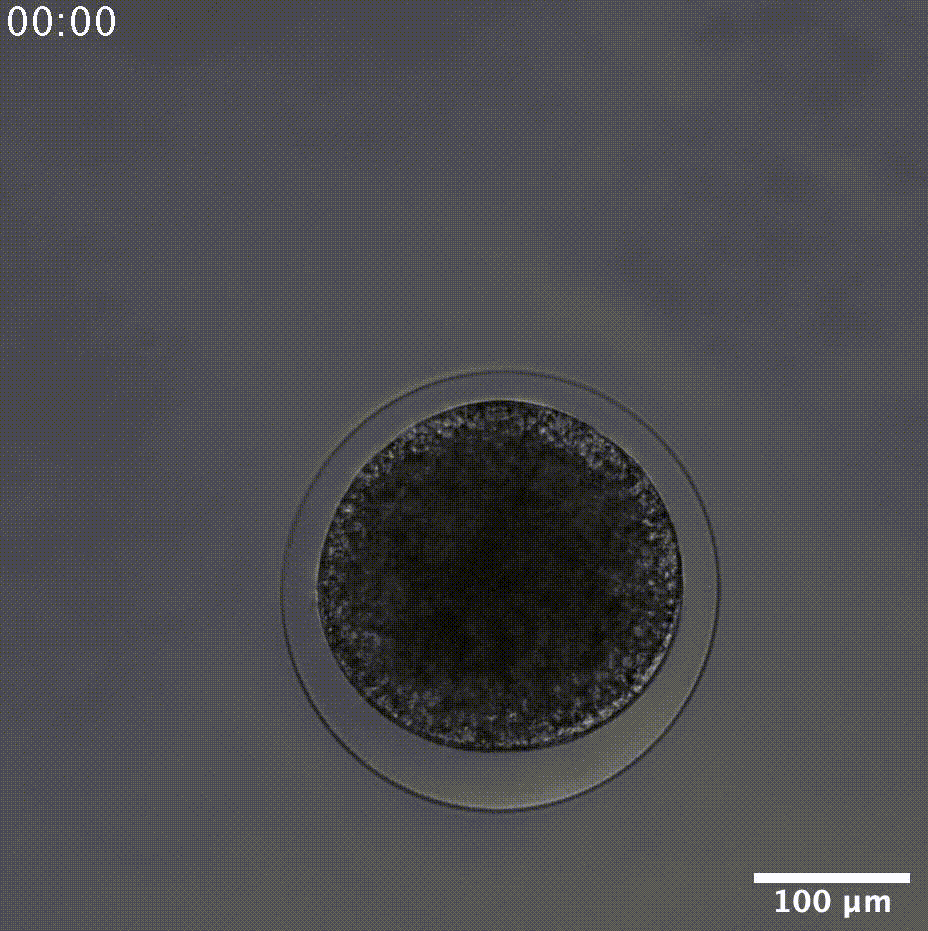

### Movie 2

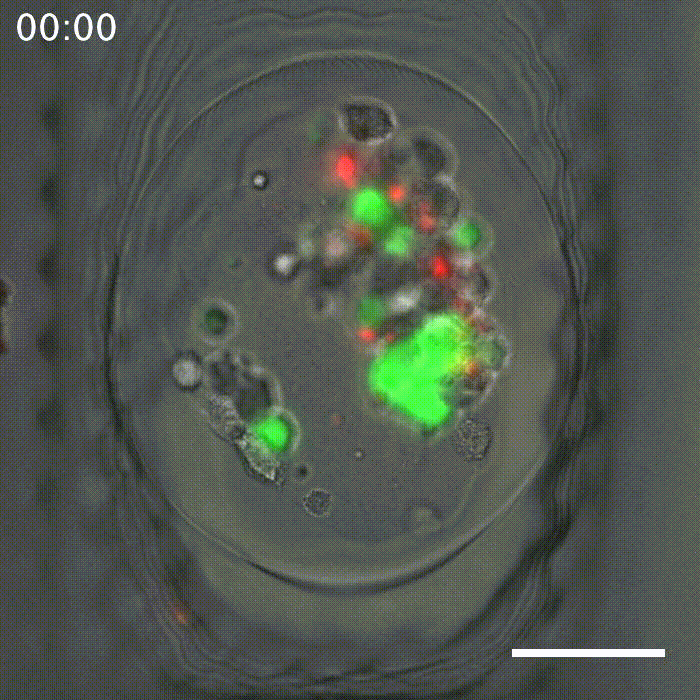

### Movie 3

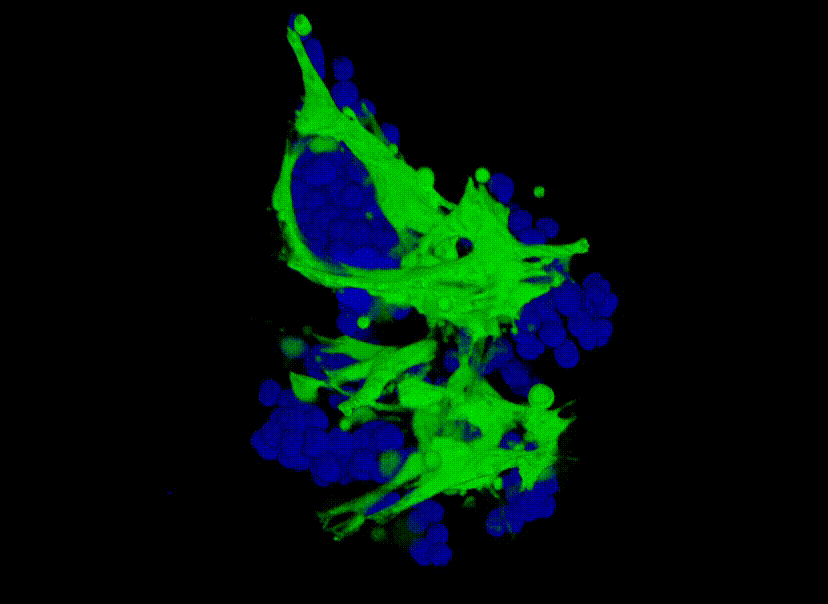
